# Mechanisms of resilience to autosomal dominant Alzheimer’s disease via oligogenic modulation of rare variants in the entorhinal cortex

**DOI:** 10.64898/2026.08.03.742644

**Authors:** Yassir El-Amri, Nelson Villalba-Moreno, Julia Urban, Maider Alzueta-Torrecillas, Cesar A Valdez-Gaxiola, Juliana Gonzalez-Perez, Zhiyuan Song, Rui Tang, Duvan Cardona-Madrigal, Andres Villegas, Susanne Krasemann, Markus Glatzel, David Aguillon, Asgeir Kobro-Flatmoen, Menno P Witter, Victoria Fernández, Haiqing Zhao, Agustín Ruiz, Claudia Marino, Diego Sepulveda-Falla

## Abstract

Two *PSEN1 E280A* carriers have presented extreme protection against autosomal dominant Alzheimer’s disease (ADAD), with over two decades of delay for dementia onset. One of them, a male heterozygous for the *RELN-COLBOS* protective variant showed increased neuronal density in the entorhinal cortex ^1^. We conducted a deep phenotyping and genotyping study of the entorhinal cortex in protected and unprotected *PSEN1* E280A cases, sporadic AD, and non-demented controls. We used single nuclei and spatial transcriptomics, whole genome sequencing, and candidate genotype-associated expression changes (GAEC) analysis. Our results showed unique neuronal and oligodendrocytic populations in the male *RELN-COLBOS* patient. Unique RELN positive inhibitory interneurons were enriched in cortical Layer I, while unique abundant ADAMTSL1 positive excitatory neurons were distributed in Layers II/III and Layer Va. These neurons and mature myelinated oligodendrocytes benefitted from increased expression of LRP6 receptor, functioning as a non-canonical receptor for the mutated Reelin protein encoded by *RELN-COLBOS*. Finally, GAEC and pathway enrichment analyses suggested that other mutations enhanced *RELN-COLBOS* effects in oligodendrocytes in the male *RELN-COLBOS* patient, explaining the phenotypic differences with his sister, a *RELN-COLBOS* carrier with no evident protection from ADAD. Our findings suggest that extreme deviations of the *PSEN1 E280A* phenotype are more likely attributed to oligogenic effects, including simultaneous mutations occurring in genes including ITGA2, involved in single molecular pathways, such as the Integrins / Focal Adhesion pathway, as potential disease modifiers for Alzheimer’s disease (AD).

## INTRODUCTION

Autosomal Dominant Alzheimer’s disease (ADAD) affects around 1% of all AD cases. Mutations in the *PSEN1*, *PSEN2*, and *APP* genes are considered causative for ADAD. Clinical and pathological presentation of ADAD is generally more severe compared to sporadic cases, with disease onset mostly in the fifth decade of life as a prominent clinical feature. The Colombian PSEN1 E280A kindred are one of the largest cohorts with ADAD, thus constituting a unique resource for the study of the pathobiology of disease and neurodegeneration in ADAD ^2,3^. Two unique outstandingly protected cases have been reported in this large population, a homozygous female APOE Christchurch (hoAPOECh) carrier with a resistant phenotype against tau pathology ^4^, and a heterozygous male RELN-COLBOS carrier showing a resilient phenotype, characterized by increased entorhinal cortex (EC) neuronal density ^1,5^. On the other hand, his sister, also carrying both the PSEN1 E280A and the RELN-COLBOS mutations, did not present with either clinical or pathological protection.

In our initial neuropathological report on the homozygous APOECh case, we used (i) single nuclei transcriptomics in frontal cortex to identify molecular changes associated with the protection against tau pathology in frontal cortex and, (ii) in parallel, we identified transcriptomic profiles associated with APOE expression in frontal cortex, occipital cortex, and hippocampus. We identified a dose-dependent effect of the APOECh mutation influencing astrocyte and microglia reactivity and homeostasis ^4^. One of our findings was the presence of RORB-positive cells in protected areas, confirming their identity as vulnerable cells in AD ^6^. Single nucleus transcriptomics have revealed molecularly distinct types of brain cells which are altered or vulnerable during the progression of the AD pathophysiological process. Early studies identified vulnerability of excitatory neurons and disease-associated microglia and astrocytes in the EC and superior temporal gyrus ^6–9^. However, single cell transcriptomic signatures do not reveal the location of cells or their cellular environment. Thus, spatial transcriptomics have become a critical tool to match molecular fingerprints with anatomic distribution, facilitating functional analysis derived from changes in gene expression ^10^. Finally, transcriptomic data in conjunction with genomic data can allow for direct interpretation of the effect of genetic variants in molecular or cellular functions by conducting association analyses between gene variants and their gene expression ^11^.

In the current study, we designed a multiomic strategy to conduct a deep phenotypic analysis on the EC of the two outstanding protected cases so far reported in the PSEN1 E280A population. We chose this brain region because of its relevance for AD ^12,13^, and because it presented with high neuronal density as the only distinctive feature in the neuropathological study of the RELN-COLBOS case ^1^. We identified unique neuronal populations benefiting from the localized effect of the mutated Reelin via a non-canonical binding to LRP6. In addition, with GAEC analyses and pathway analysis of genomic data, we identified additional genetic variants associated with distinctive oligodendrocyte populations and the probable modulation of myelinization by the Integrin signaling pathway. Our findings suggest that even though the RELN-COLBOS mutation does have a direct effect on neuronal and non-neuronal populations, it is possible that the cumulative effect of additional mutations generated the extremely protected phenotype observed in this ADAD patient.

## Methods

### Human brain tissue samples

Frozen and formalin fixed paraffinized samples were supplied by the Brain Bank of the Neurosciences Group of the University of Antioquia, Medellin, Colombia. Brains were collected after signature of informed consent by the patients or their family members at the time of brain donation. Donation procedures and informed consent were approved by the Ethical Committee of the Faculty of Medicine at the University of Antioquia. All samples used in this study had a postmortem interval of less than 12 hours. Selection of cases for each experimental procedure was based on sample availability at the time of the experiments; their basic clinical profiles and demographic characteristics of the cases are summarized in Supplementary Table 1. Hippocampal samples from both hemispheres were used, one snap frozen at the time of donation and the contralateral sample fixed in formalin (Fig. 1A).

**Figure 1.**
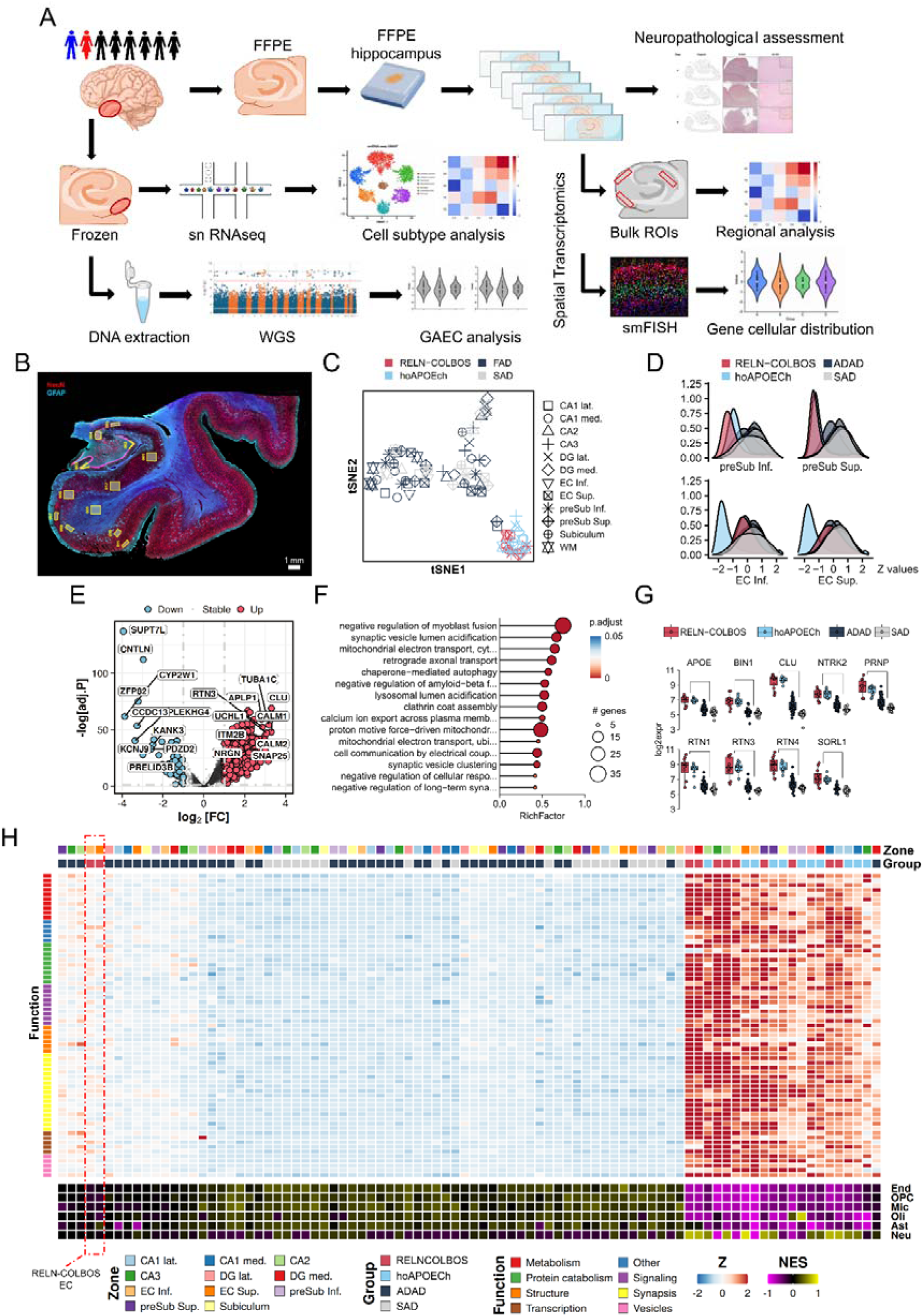
Bulk spatial transcriptomics analysis of hippocampal areas in protected and unprotected AD cases. A. Study design and data analysis overview. Postmortem hippocampi or specifically EC from PSEN1 E280A protected cases (blue male, heterozygous RELN-COLBOS case; red female, hoAPOECh case), and PSEN1 E280A unprotected cases and controls (black) were analyzed. FFPE tissue was used for neuropathological assessment and spatial transcriptomics, while frozen entorhinal cortices were used for snRNA seq and whole genome sequencing. B. Immunofluorescent microphotograph of the whole hippocampus from the male RELN-COLBOS case, stained with NeuN (Red), and GFAP (Cyan), the regions of interest selected for bulk spatial transcriptomic analysis are depicted in yellow shapes. C. tSNE dimensionality reduction across all cases and hippocampal areas (8 samples, 96 ROIs). DG: dentate gyrus, EC: entorhinal cortex, preSub: presubiculum, WM: white matter, lat.: lateral, med.: medial, Inf.: inferior, sup.: superior. D. Density plots of individual gene expression across all genes in the presubiculum and EC. E. Volcano plot of differentially expressed genes (DEGs) between protected and non-protected cases, all regions combined. F. Lollipop plot of enriched GO Biological Process terms among DEGs between protected and non-protected cases, all regions combined. G. Boxplot for GO:1902430 (negative regulation of amyloid-beta formation) genes expression across all ROIs, by condition. H. Heatmap of genes commonly dysregulated in protected vs. ADAD cases, by area. Color codes indicate patient, hippocampal, functional and molecular contribution, together with deconvoluted cell type contribution,

### GeoMx Digital Spatial Profiler (DSP) analysis

Nanostring GeoMX Digital Spatial Profiler (DSP) was used to obtain human whole transcriptome data by counting uniquely indexed transcripts for over 18,000 protein-coding genes. Formalin-fixed paraffin-embedded (FFPE) postmortem human brain tissue samples of the hippocampal structures were dissected, including the male RELN-COLBOS carrier, the hoAPOECh carrier, four PSEN1 E280A ADAD cases and two sporadic AD controls. Formalin-fixed hippocampal samples were mainly taken from posterior hippocampi. The sections were mounted on Superfrost slides (Thermo Fisher) and were prepared and processed according to manufacturer instructions (user manual MAN-10150-02, nanostring.com). Slides were stained with GFAP and NeuN antibodies and scanned at 20x magnification to help guide region of interest (ROI) selection. The following 12 different anatomically relevant cortical compartments (zones) were delimited: lateral CA1 (CA1 lat.), medial CA1 (CA1 med.), CA2, CA3, lateral Dental Gyrus (DG lat.), medial Dental Gyrus (DG med.), inferior EC (EC inf.), superior EC (EC sup.), inferior presubiculum (preSub. inf.), superior presubiculum (preSub. sup.), subiculum and White Matter (WM). Altogether, 96 ROI were collected and one No Template Control (NTC) to detect contamination in the library preparation. Library preparation, QC and sequencing were performed according to the GeoMx® DSP NGS Readout User Manual, MAN-10153-02 (nanostring.com). The FASTQ files generated by the sequencing run were processed into digital count conversion (DCC) files using the NanoString GeoMx NGS Pipeline which were then used for data analysis.

### GeoMX Data analysis

All analyses were conducted using R (version 4.4; R Foundation for Statistical Computing, Vienna, Austria) ^14^. Data quality control (QC) and preprocessing followed the GeoMX Workflows guidelines ^15^, incorporating metrics such as minimum segment reads, percentage of stitched and aligned reads, sequencing saturation, and performance of negative control probes. Samples that did not meet QC thresholds and genes detected in fewer than 10% of ROI were excluded. Quartile 3 (Q3) normalization was applied to the filtered expression data. To explore large-scale transcriptional variability across samples, unsupervised dimensionality reduction was performed using t-distributed stochastic neighbor embedding (t-SNE).

A single experiment without technical replicates was conducted given tissue availability limitations. Differential expression analysis was first conducted between protected and both ADAD and SAD cases for all regions using edgeR package ^16^. A functional enrichment analysis was conducted with the resulting dysregulate genes (adjusted p value < 0.05, |log (fold change) | ≥L1 (Supplementary Table 2). Subsequently, individual differential expression analyses were run for each protected case against the AD cases within each selected hippocampal region. Genes that were consistently dysregulated in at least 8 out of the 12 regions (≥75%) were visualized in a heatmap and functionally annotated based on known biological roles. Whole transcriptomics expression data was used to estimate the relative proportions of specific brain cell types using the BRETIGEA package ^17^. Functional enrichment analysis of Gene Ontology (GO) terms was conducted on the dysregulated genes using the clusterProfiler package ^18^. Redundant GO terms were further filtered based on hierarchy and gene overlapping similarity, therefore keeping the most specific terms. All visualizations were generated using ggplot2 package ^19^.

### Single-nucleus RNA sequencing

Snap-frozen EC samples, predominantly from anterior hippocampi from the male RELN-COLBOS case, the female RELN-COLBOS case, the hoAPOECh case, seven ADAD cases, and three controls were individually dissociated, and nuclei were isolated using the Nuclei EZ Prep kit (Sigma, #NUC101). Briefly, 50 mg of frozen tissue was homogenized in a precooled douncer with 1 mL of ice-cold Nuclei EZ Isolation Buffer. After homogenization, 3 mL of additional buffer and 2 µL of Collagenase II (0.05U) were added to the suspension and incubated on ice for 5 min. The nuclei suspension was centrifuged at 500 x g for 10 min at 4°C, and the pellet was resuspended in 4 mL of cold Nuclei Suspension Buffer and filtered through a 30 µm cell strainer. The filtered suspension was centrifuged again at 500 x g for 10 min at 4°C, and the resulting pellet was washed with 4 mL of ice-cold 2% BSA in PBS. A second filtration was performed through a 30 µm cell strainer, followed by another centrifugation (500 x g, 5 min and 4°C). The resulting pellet was resuspended in 100 µL ice-cold 2% BSA in PBS. Nuclei were stained with Trypan Blue stain 0.4% (cat. #T10282, InvitrogenTM, Carlasbad, CA, USA) and counted using a Neubauer chamber. A final concentration of 1,000 nuclei/µL was prepared for loading onto the Chromium Controller (10X Genomics). Library construction was performed using the Chromium Single Cell 3’ Library & Gel Bead Kit v3.1 (10X Genomics) according to user’s guidelines and sequenced on the NextSeq 4000 (Illumina Inc., San Diego, CA, USA) at the Next generation sequencing facility at the University of Kiel, Germany.

Sequencing reads were mapped to the human reference transcriptome (GRCh38) - 2024-A using the 10x Genomics Cell Ranger pipeline (version 7.2.0) with default parameters. Downstream analysis was performed using Seurat 4.0 in R 4.4. Cells with fewer than 200 detected genes and with more than 5% of reads mapped to mitochondrial and ribosomal genes were filtered out. Genes expressed in less than 10% of cells were removed. Doublets were identified using the DoubletFinder package ^20^ and removed, assuming a doublet rate formation of 7%.

All samples were merged into a single Seurat object. Data were then normalized and scaled using the SCTransform function in Seurat using the default parameters and regressing the percentage of mitochondria. To integrate all samples, principal component analysis (PCA) was performed on the normalized data, and batch effects across samples were corrected using Harmony on the PCA embeddings. Non-linear dimensionality reduction was performed by running UMAP on the first 40 PCs. Cell clustering was performed using the *FindNeighbors* function with the first 40 PCs as the input, followed by *FindClusters* with a resolution of 1. To find cluster-specific markers, we defined cell type specific gene sets for the major cell types (neurons, astrocytes, oligodendrocytes, oligodendrocyte precursor cells (OPC), microglia and endothelial cells) using the Allen Brain Atlas and mapped them to the obtained clusters. After assigning major cell type clusters, we subsetted the data of each cluster, reprocessed and re-clustered the cells to obtain a higher-resolution identification of cellular subpopulations within the same cell type.

### Whole genome sequencing

Brain tissue samples (20 mg) were homogenized in MagMAX™ DNA Cell and Tissue Extraction Buffer (#A32702, Thermo Fisher Scientific) using a rotor-stator homogenizer (Tissue Ruptor II; Qiagen). DNA extraction was subsequently performed using the MagMAX™ DNA Multi-Sample Ultra 2.0 Kit (#A36570, Thermo Fisher Scientific) according to the manufacturer’s instructions. DNA quantity and quality were assessed using the Agilent TapeStation system (Agilent Technologies Inc., Santa Clara, CA, USA). Genomic DNA samples were subjected to quality control prior to library preparation. Whole-genome sequencing libraries were prepared using the Ultima TruSeq PCR Plus library preparation kit (Illumina-compatible chemistry adapted for the Ultima platform). Sequencing was performed by Macrogen Inc. (Seoul, Republic of Korea) on the Ultima UG100sequencing platform, targeting approximately 90 Gb of sequence data per sample, corresponding to an average genome coverage of approximately 30×. Raw sequencing data were delivered as CRAM files following standard quality control procedures.

Variant calling was performed using Efficient DV, an analysis pipeline adapted from DeepVariant ^21^ for Ultima Genomics sequencing data. The pipeline operates on aligned CRAM files through three sequential stages for each sample. First, active genomic regions are identified, local haplotype assembly is performed, reads are realigned, and candidate variants are defined (*make_examples*). Second, a TensorRT-based deep learning model processes the candidate variants to compute genotype likelihoods and quality scores (*call_variants*). Third, multi-allelic records and indel-overlapping variants are resolved; variants are annotated with type, cycle-skip status, and genomic interval membership, filtered by quality thresholds, and written to a gVCF output file (*post_process*).

The individual gVCF files were further joined using GLNexus ^22^ to generate a cohort-level VCF. Variant quality control was then performed using bcftools. Variant sites with a Phred-scaled quality score (QUAL) <20 were excluded, and genotype calls with a Genotype Quality (GQ) score < 20 were converted to missing genotypes. The resulting VCF was normalized against the GRCh38 reference genome using bcftools norm, including reference-based allele normalization to ensure a consistent representation of variants across samples. Functional annotation of the variants was subsequently performed using ANNOVAR ^23^. After annotation, variants with a missing genotype rate >10% across samples were excluded. Finally, to perform a rare variant candidate search, gene variants were then further filtered to retain only those that were considered as rare, defined as having an allele frequency below 1% in the gnomAD v4.1 genome database (general allelic frequency, gnomad41_genome_AF ≤ 0.01) ^24^.

To find genotype-association expression changes (GAEC) between the gene variants of the studied cases and their expression in the major cell types, we conducted a single-cell-resolution Poisson mixed-effects (PME) regression. The UMI counts of a gene in the single nuclei were modelled as a function of genotype, adjusting for confounders (log(UMI count), gene expression PCA, post-mortem interval (PMI)). Random-effect covariates account for donors. The model’s significance was evaluated using a likelihood ratio test (LRT). Common and distinctive eGenes (genes which expression is significantly altered by the genotype) were plotted using ComplexUpset ^25^.

### Variant pathogenicity prediction

Given that the carriers of the protective variants APOECh and RELN-COLBOS were also carriers of other rare mutations in other genes, we conducted gene enrichment analysis for cellular pathways that would be simultaneously affected by several exonic deleterious variants, that would affect directly protein-protein interactions. The pathogenic potential of each missense variant belonging to the male RELN-COLBOS case and identified in the genes included in the KEGG pathways hsa04510 and hsa04512, was assessed using three complementary computational predictors: AlphaMissense ^26^, Combined Annotation Dependent Depletion (CADD) ^27^, and PrimateAI-3D ^28^. Variants were specified at the protein level and mapped to canonical human UniProtKB sequences for sequence-based methods and to GRCh38 (hg38) genomic coordinates for variant-effect scoring.

Missense pathogenicity was first scored using precomputed AlphaMissense predictions for the human proteome (hg38 release). Each variant was matched to its canonical UniProtKB entry and protein-level substitution (e.g., COL6A6, UniProtKB A6NMZ7; FN1, UniProtKB P02751). For FN1, residue numbering was offset by +91 to reconcile our transcript, which lacks the alternatively spliced extra-domain B (EDB), with the canonical UniProtKB P02751 sequence. AlphaMissense outputs a continuous pathogenicity score in the range [0, 1]; variants were classified using the published thresholds of <0.34 (likely benign), 0.34–0.564 (ambiguous), and >0.564 (likely pathogenic).

Genome-wide deleteriousness was then evaluated with the CADD framework, version 1.7 (GRCh38). Variants were submitted as a minimal VCF (chromosome, position, reference, alternate allele) to the CADD web server, and the scaled PHRED-like score was retained for the non-synonymous coding consequence of each variant. The PHRED score represents the rank of a variant relative to all ∼8.6 billion possible single-nucleotide variants in the genome, on a −10·log_₁₀_ scale; a score of 10, 20, or 30 corresponds to a variant ranking in the top 10%, top 1%, or top 0.1% most deleterious substitutions, respectively ^27^. Missense pathogenicity was last evaluated using precomputed PrimateAI-3D predictions. Each variant was matched to its corresponding RefSeq entry and protein-level substitution. For RELN and FN1, residue numbering was offset by −2 and +6, respectively, to reconcile canonical UniProtKB positions with the specific RefSeq transcript isoforms indexed by the database. PrimateAI-3D outputs a continuous pathogenicity score in the range [0, 1] based on a deep learning model of 3D protein structure and cross-species evolutionary conservation, where higher values indicate a greater likelihood of a functionally deleterious effect; variants were classified as benign (score < 0.821) or pathogenic (score ≥ 0.821).

### Rare deleterious exonic variants burden analysis pipeline

To locate individual mutation carriers within the population distribution of pathway variant burden, we used the high-coverage 1000 Genomes Project (1000G) as reference. The dataset was first processed to retain only the 2504 unrelated subjects and high-quality biallelic SNVs, which were then annotated with NCBI Reference Sequence (RefSeq), gnomAD v4.1, and CADD information. We next kept the rare SNVs (gnomAD v4.1 genome AF < 0.01) falling in genes of the pathways of interest. These variants were grouped into missense and high-confidence functional-disrupting categories: missense variants qualified at CADD_PHRED ≥ 20, whereas high-confidence disrupting SNVs (e.g. stop-gain) were retained regardless of CADD. Burden was then calculated per subject as the number of qualifying pathway alleles carried, each counted once irrespective of heterozygous or homozygous dosage. For each case and pathway, we counted reference subjects with lower, equal, and equal-or-higher burden; the empirical percentile used the mid-rank position (subjects below + ½ subjects equal), and the upper-tail probability was the fraction of reference subjects with burden greater than or equal to the case value.

The RELN-COLBOS male and female carriers and the hoAPOECh carrier were projected against the population, applying the same criteria used for the 1000G samples. Three pathways were evaluated: hsa04141 (control: 109 of 112 genes mapped), hsa04512 (84 of 86), and hsa04510 (141 of 151), with 2097, 4613, and 5692 qualifying reference variants, respectively. The low case burden in hsa04141 reflected genotype absence rather than a mapping or filtering error.

The analysis was implemented using the pathway_burden_pipeline github repository (https://github.com/neurogeneticafundacioace-source).

We further performed KEGG enrichment analysis for each 2504 unrelated samples from 1000G, using the genes whose variants met the criteria described above. The resulting pathways were ranked by their frequency of appearance across the full sample set, and we assessed whether these pathways were likewise enriched in the EC study samples.

### Combinatorial single molecule Fluorescence in-situ hybridization (smFISH)

The EC region, anatomically defined from hematoxylin and eosin staining, were dissected from ten microns thick FFPE hippocampal slice cuts from the male RELN-COLBOS case, the hoAPOECh case, two sporadic AD cases and one healthy control. The dissected ECs were placed on a Superfrost slide sent for DAPI staining and multiplex smFISH processing at the manufacturers side (Resolve BioSciences GmbH, Monheim am Rhein, Germany) as previously described ^29^, using a gene panel (Suppl. Table 5) designed from the cellular clusters identified by snRNAseq. QuPath (v.0.3.0) ^30^ was used to segment cells based on their DAPI images, then used Fiji (v.1.52n) ^31^ and the Molecular Cartography plug-in (Resolve Biosciences) to count genes in each cell. For the DAPI image, we also manually annotated different cortical layers using different cortical layers markers (LAMP5, RORB, ADAMTSL1). The cell-gene count matrix was then input into Seurat (v.3.2.3) ^32^ for downstream analysis. For each of the 5 samples of Resolve spatial data, cells with gene counts below the 15th percentile were excluded from the analysis. Counts data were then normalized using NormalizeData with the default LogNormalize method. Afterwards, normalized counts were scaled and centered using the ScaleData function. Gene expression correlation analysis was performed to identify sets of co-expressed genes. Their expression levels were quantified within annotated cortical layers to evaluate layer-specific differences, and their spatial distributions were visualized across tissue sections.

For specific combination of markers, the proportion of positive cells for each combination was quantified as the number of cells that contain that combination divided by the total number of cells per layer. Cells with RELN transcripts within each sample were annotated as RELN-positive cells. For the neighborhood analysis, fixed-radios nearest neighbors (FRNN) were computed using a radius of 50 around the RELN-positive cells using the frNN() function from the dbscan R package ^33^. For each RELN-positive cell, the expressions of the low-density lipoprotein receptor-related proteins LRP4, LRP6 and LRP8 were quantified. A cell was considered to express one of these genes if its count was greater than zero. Only RELN-positive cells containing at least three neighboring cells were retained. To normalize for gene-specific abundance differences, the aggregated neighbor expression for each receptor was divided by the number of neighboring cells and further scaled by the total fraction of cells in the sample expressing that receptor, yielding a relative expression enrichment score per receptor and RELN-positive cell.

### Golgi-Cox staining and Immunofluorescence

Coronal slices from 6 months old mice, either RELN-COLBOS transgenic (n=4) or wild type (n=3), were prepared for Golgi-Cox staining using the FD Rapid Golgi Stain Kit (PK401, Hölzel Diagnostika Handels GmbH, Cologne, Germany) according to manufacturer instructions. Full cortical images were assembled from 20x magnified pictures taken with a Leica TCS SP8 confocal laser scanning microscope (Leica Microsystems, Mannheim, Germany) using the brightfield mode. For immunofluorescence imaging from human hippocampi and murine FFPE tissue, 4 microns thick sections were mounted in Superfrost plus slides, deparaffinized, and heat-induced epitope retrieval was performed using R-Universal buffer (AP0530-500; Aptum Biologics, Southampton, UK) in a pressure cooker for 20 min, sections were incubated for 10 min in 0.25% X-100 Triton in PBS and were then blocked for 1 h with blocking medium (MAXblock_^TM^, #102224, Active Motif Inc., Waterloo, Belgium) followed by incubation with primary antibodies dissolved in MAXbindTM Staining Medium (#102227, Active Motif Inc., Waterloo, Belgium) at 4°C overnight. Primary antibodies used were anti-Reelin (Reelin E-5, 1:100, sc-25346, Santa Cruz Biotechnology, Inc. Heidelberg, Germany), anti-Hexaribonucleotide Binding Protein-3 (NeuN, 1:200, #26975-1-AP, Protein Tech Germany, Planegg-Martinsried, Germany), and anti-Opalin (Opalin, 1:400, AMAb91685, Atlas antibodies AB, Stockholm, Sweden). After washing the slides with MAXwashTM Washing Medium (#102224, Active Motif Inc., Waterloo, Belgium), specific binding detection was conducted with secondary antibodies incubated at room temperature for 1 h: AF488 (Donkey-a-Ms, 1:200, #A21202), AF555 (Goat-a-Rb, 1:200, #A21429), (#MA5-16785 FITC Ms IgG1, #A21241 AF647 MsIgG2b; 1:200). To avoid autofluorescence and lipofuscin signal, the slides were incubated for 3-4 min in True Black® Lipofuscin Autofluorescence Quencher, 20X (23007, Biotium, Fremont, CA, USA) diluted 20x in 70% ethanol. After washing, mounting was performed with 4′,6-Diamidino-2-phenylindole (DAPI) Fluoromount-G® (#0100-20, SouthernBiotech, Birmingham, AL, USA) for nuclear counterstaining. High-resolution images were obtained with a Leica TCS SP8 confocal laser scanning microscope (Leica Microsystems, Mannheim, Germany) using a 20X immersion lens objective.

### Protein-protein interaction modeling

AlphaFold3 was used to predict putative receptor-ligand interfaces for hypothesis generation. It was used to predict the joint 3D structures of the following proteins and modified peptides ^34^: RELN (UniProtKB: P78509), LRP4 (UniProtKB: O75096), LRP6 (UniProtKB: O75581), WNT3a (UniProtKB: P56704), ITGA2 (UniProtKB: P17301), C Terminal Region (CTR)-RELN WT (amino acid sequence: LVSTRKQNYMMNFSRQHGLRHFYNRRRRSLRRYP), and CTR-RELN COLBOS (amino acid sequence: LVSTRKQNYMMNFSRQHGLRRFYNRRRRSLRRYP). Protein complexes were run at a 1:1 stoichiometry using the hosted AlphaFold Server (https://alphafoldserver.com, accessed between February 2025 and June 2026). Obtained models were subsequently imported into UCSF ChimeraX software (Ver. 1.10) ^35^ to identify and characterize surface cavities with the “KVFinder algorithm” ^36,37^.

In parallel experiments, Protein-Protein Interactions (PPIs) were screened using the structure-based genome-wide PPI database PrePPI ^38^. Predicted interactions and their structural templates were manually inspected and analyzed using PyMOL v2.5.2 [Schrodinger, L. (2010) The PyMOL Molecular Graphics System, Version v2.5.2]. Independent structural models were generated using AlphaFold3 ^34^ and used to validate the predicted interaction interfaces. Comparative analyses of LRP4, LRP6, and LRP8 domain architectures were performed using the UniProt Family & Domains annotation resources ^39^. Full-length protein structures predicted by AlphaFold were further examined in PyMOL v2.5.2 to assess structural conservation and differences among family members.

### ELISA binding assays

High-binding 96-well plates were coated with 100 μL per well of biotinylated Human DKK1 Fc-tag/Avi-Tag (Acro Biosystems, cat. #DK1-H82F5), C-terminal region (CTR)-RELN-Wild Type (WT) 184I Fc-tag (Innovagen, lot. #16577.04), or CTR-RELN-COLBOS 184J Fc-tag (Innovagen, lot. #16578.04) at a final concentration of 5 μg/mL. Both CTR-peptides were previously designed and validated by the Arboleda-Velasquez laboratory ^1^. All coating solutions were prepared in coating buffer (Candor Bioscience, cat. #121125), and plates were incubated overnight at 4 °C on a shaker to ensure uniform protein adsorption. Following incubation, plates were washed four times with 300 μL per well with washing buffer consisting of 0.05% Tween-20 in TBS 1X (pH 7.4, Thermo Fisher, cat. 28360). Wells were then blocked with 300 μL per well Blocking Buffer (2% BSA in washing buffer) for 1.5 hours at 37 °C without shaking. After an additional washing step, 100 μL per well of recombinant human LRP6 Fc Chimera (R&D Systems, cat. #1505-LR), diluted in sample dilution buffer (0.5% BSA in washing buffer, pH 7.4), was added and incubated at 37 °C for 1 hour to allow ligand-receptor binding. Plates were then washed and incubated for 1 hour at 37 °C with mouse anti-LRP6-HRP detection antibody (100 μL per well, Novus Biologicals, cat. #FAB1505H), diluted 1:2,000 in sample dilution buffer. After washing, substrate development was performed by adding 200 μL per well of substrate solution (R&D Systems, cat. #899517.01) and incubating plates for 15 minutes at 37 °C in the dark. The reaction was stopped with 50 μL per well Stop Solution (R&D Systems, cat. #895926.02), and absorbance recorded at 450 nm using a microplate reader (Cytation 5, Agilent BioTek). Data was then analyzed and plotted to obtain the area under the curve (AUC) and the nonlinear relationship between logarithmic concentrations and normalized percentage of LRP6 binding using GraphPad Prism 11. Statistical differences between AUC were analyzed on two independent experiments (n=3 technical replicates) using one way ANOVA followed by Tukey’s test for multiple comparisons. Any p value less than 0.05 was considered statistically significant.

## RESULTS

### Distinctive spatial transcriptomic profile of the EC of the male PSEN1 E280A / RELN-COLBOS patient

Hippocampus, and more specifically, the EC is a brain region that has shown a degree of protection in the two extremely protected cases ADAD with PSEN1 E280A ^1,4^. In fact, the male RELN-COLBOS case showed decreased tau pathology detected by PET and increased density of superior layers of the EC ^1^. This localized protection can potentially be associated with specific molecular pathways in specific cell populations. To explore the transcriptomic profile of hippocampal areas in ADAD protected cases, a GeoMx Digital Spatial Profiler (DSP) analysis was performed on human brain FFFP from 8 patients, including both protected cases (RELN-COLBOS and hoAPOECh cases), ADAD cases and sporadic cases. Twelve different hippocampal areas were identified in all cases as regions of interest (ROI) for spatial bulk transcriptomic analysis (Fig. 1b). Interestingly, the dimensionality reduction using tSNE showed a clear segregation between protected and non-protected cases among all areas, regardless AD etiology or age, except from two isolated ROIs from two unprotected cases (Fig. 1c). This clear difference was also evident in density plots for all ROIs, except for the EC, in which only the hoAPOECh showed a distinctive profile (Fig. 1d, Suppl. Fig. 1). Next, we directly compared protected and non-protected cases combining all hippocampal areas, resulting in 1,027 significantly differentially expressed genes (Fig, 1e). Functional enrichment analyses of these genes showed upregulation of synaptic-related functions, mitochondrial function, autophagy and lysosomal function, and negative regulation of amyloid beta formation (Fig. 1f, Suppl. Table 2). Interestingly, for the latter function, genes included *APOE, CLU, PRNP,* and *SORL1* (Fig. 1g). In fact, a subset of 67 genes was commonly upregulated in both protected cases in most hippocampal areas, except for both entorhinal ROIs in the RELN-COLBOS case. These genes are involved in metabolism, synaptic function, and protein catabolism, among others ^40^. Deconvolution analysis of the whole bulk transcriptome data indicated a high neuronal contribution to the upregulated expression profile (Fig. 1h, Suppl. Table 3).

### Increased association between rare gene variants and single nuclei gene expression in the male PSEN1 E280A / RELN-COLBOS patient

Given that the EC from the RELN-COLBOS male case showed higher neuronal density (Suppl. Fig. 2) and a distinctive bulk transcriptomic profile, we isolated nuclei from frozen brain material for single nuclei RNA sequencing (snRNA-seq). We included frozen EC tissue from the RELN-COLBOS sister, together with the hoAPOECh case, seven other PSEN1 E280A cases, and three controls (Suppl. Table 1). The RELN-COLBOS female carrier did not present a strong protective phenotype as evidenced by earlier age of onset and low neuronal density in the EC (Suppl. Fig. 2). 103,379 nuclei were successfully filtered for analysis, encompassing six major cell types (Fig. 2a). From those cells, neurons represented the highest percentage (50.52%), followed by astrocytes and oligodendrocytes (20.54% and 13.7% respectively). Control patients showed a relatively higher contribution in neurons, while the female RELN-COLBOS patient contributed with a relatively higher number of cells among oligodendrocytes and endothelial cells. Both the male RELN-COLBOS case and the hoAPOECh case showed similar cell type distribution to the other ADAD cases (Fig. 2b). In parallel, we conducted Whole Genome Sequencing (WGS) of these cases to evaluate the possible influence of rare genetic variants detected in the three protected cases in the corresponding gene expression, by using GAEC analysis. We identified 775 eGenes , some of them commonly associated with more than one cell type (Fig. 2c). Both protective gene mutations, RELN-COLBOS and APOECh, have an allelic frequency below 0.01. Both protected cases carrying these mutations carry also a high number of equally rare variants (70644 for male RELN-COLBOS carrier, and 50898 for the hoAPOECh case). Additionally, the female RELN-COLBOS case shows a similarly high count of rare variants (49750). The number of rare variants identified in the three cases was within the range of the number of rare variants identified in general population ^41^. Interestingly, even though rare variants of all the cases were included for the analysis, the large majority of the eGenes identified among these three cases are associated with genetic variants identified in the male RELN-COLBOS case, either alone or shared with his sister (Fig. 2d-e, Suppl. Fig. 3). Besides the RELN-COLBOS mutation, the RELN gene shows eleven additional mutations in both RELN-COLBOS carriers. However, these mutations do not associate with RELN expression in either of those cases (Fig. 2f). On the other hand, rare gene variants carried by the RELN-COLBOS cases associated with eGenes included potassium and calcium channels, and evidenced in several cell types, mainly in astrocytes, oligodendrocytes, and oligodendrocyte precursor cells (OPC) (Fig. 1g-h). When gene enrichment analysis is performed in all the eGenes identified in both RELN-COLBOS carriers associated with rare gene variants, we observe genes involved in regulation of signal transduction, second messenger mediated signaling, and signaling receptor activity. Additionally, these genes are also part of KEGG pathways including GABA, Calcium, Oxytocin, MAPK, Rap1, Ras, and cGMP-PKG signaling. Other KEGG pathways involved were axon guidance, circadian entrainment, long term depression, and serotoninergic and cholinergic synaptic pathways. These pathways were identified for eGenes associating in one or several cell types, mainly in oligodendrocytes or OPC (Fig. 2i, Suppl. Table 4).

**Figure 2.**
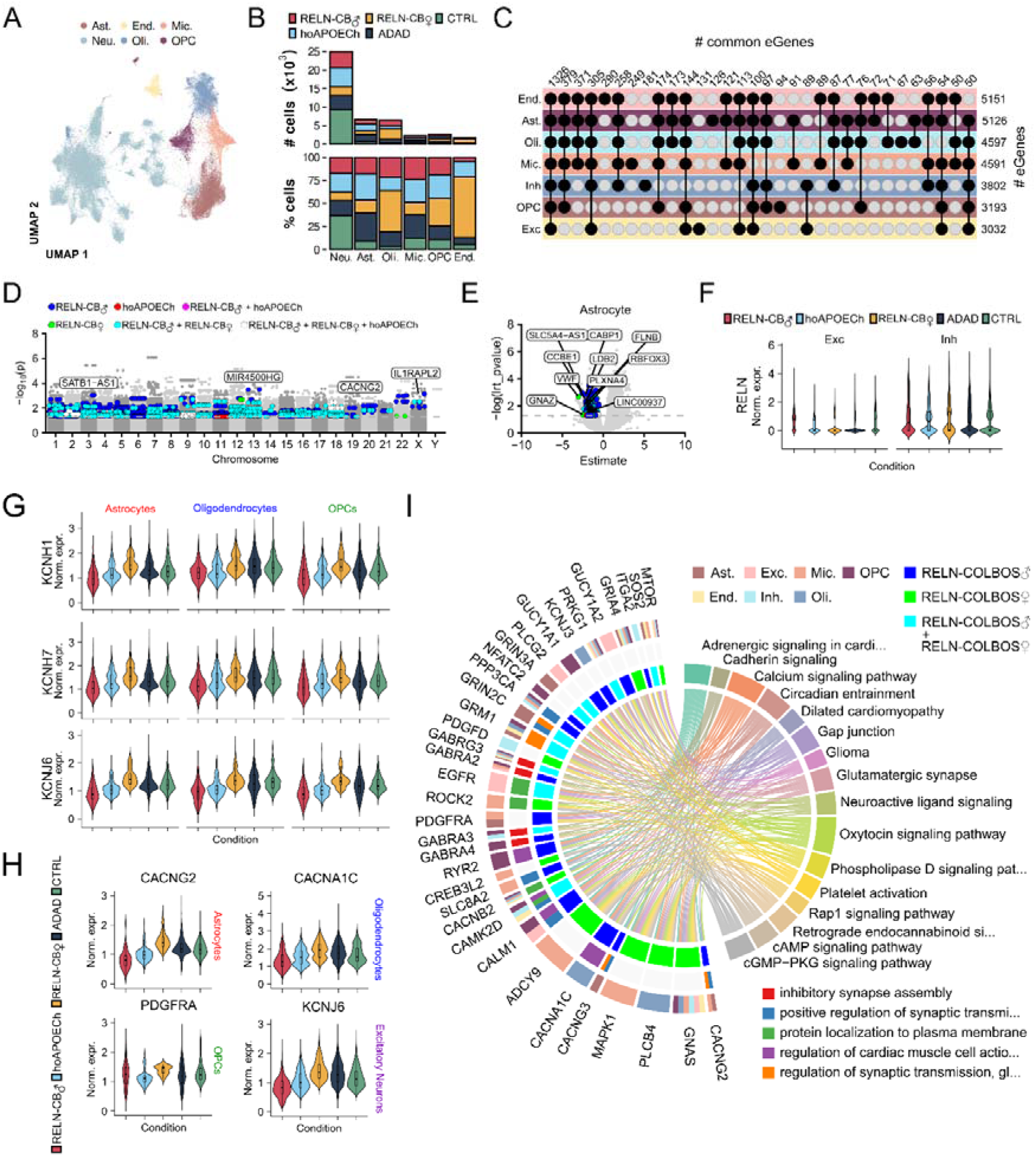
Single nuclei transcriptomics and candidate Genotype-Associated Expression Changes (GAEC) analysis in entorhinal cortices of protected and unprotected AD cases. A. Uniform manifold approximation and projection (UMAP) projection of single-nuclei transcriptome profiles from 103379 cells integrated, showing 6 major cell types. Astrocytes (Ast.) = 21232, Endothelial (End.) = 2817, Microglia (Mic.) = 6898, Neurons (Neu.) = 52225, Oligodendrocytes (Oli.) = 14162, Oligodendrocyte Precursor Cells (OPC) = 6045. B. Stacked bar plot of cell counts per cell type and condition, normalized by number of cases per condition. C. UMAP of neuronal cells, labelled as excitatory or inhibitory. C. UpSet plot of cell-type-specific vs. shared eGenes (y-axis), connected by lines: x-axis truncated below 50 eGenes per category. D. Manhattan plot of rare-variant-expression associations in astrocytes. Variants with LRT p < 0.05 are colored by group: RELN-COLBOS♂ (blue), hoAPOECh (red), or RELN-COLBOS♀ (green); gray line marks LRT p = 0.05. E. Volcano plot for significant eGenes in astrocytes according to the estimate of their effect using the GAEC analysis. F. Violin/box plots of RELN expression in excitatory and inhibitory neurons, by condition. G. Violin/box plots of inwardly rectifying potassium channel genes (linked to rare variants) in astrocytes, oligodendrocytes and OPCs. H. Violin plots of select associated genes by cell type and condition. I. Chord diagram of enriched KEGG pathways and GO terms for selected genes, labelled by cell-type association and RELN-COLBOS carrier.

### Increased number of RELN+ inhibitory Layer I neurons in the male PSEN1 E280A / RELN-COLBOS patient

We identified 19 neuronal clusters, 10 for excitatory neurons, and 9 for inhibitory neurons (Fig. 3a). Next, we identified six RELN positive (RELN+) neuronal clusters, including four clusters in inhibitory neurons (I03, I05, I07, and I08). Clusters I03 and I07 are LAMP5 positive; cluster I05 is PVALB and VIP positive. Additionally, cluster I07 was also SST positive, and cluster I08 ADAMTSL1 and VIP positive (Fig. 3b). A deeper assessment of gene expression patterns among inhibitory neurons clusters showed that RELN+ clusters are also ADARB2 positive, but each of them presented distinctive gene expression signatures. For instance, cluster I03 is also positive for DOCK5, GAD2, LAMP5, and TOX2, suggesting its identity as neurogliaform interneurons ^42^. On the other hand, I05 and I08 cells were positive for NPAS1, NPAS3, RGS12, and CXCL14, characteristic of upper layers non neurogliaform inhibitory neurons ^42^. I05 neurons distinctively expressed FREM2, while I08 neurons distinctively expressed SCML4, KLHL1, SMOC1, and GAD1. Low LAMP5 and VIP expression in I05 cells indicate a possible localization in Layer I, while SCML4 and KLHL1 in I08 cells extend their possible localization between Layers I-II (Allen Brain Atlas) ^43^. I07 neurons showed the highest expression of RELN, together with RGS12, CXCL14, NDNF, GAD2, LAMP5, and TOX2, indicating their possible location in Layer I, but leaving their identity between neurogliaform and non neurogliaform cells uncertain ^42^ (Fig. 3c). We chose RGS12 and CXCL14 as distinctive markers highly expressed among some RELN+ cells, that would associate with specific cortical Layer location and cell type, to analyze differential gene expression among them. Thus, we examined differentially top expressed genes, clarifying the gene expression signature between the different RELN+ clusters. Cluster I05 showed a more homogenous gene expression among individual cells, including expression of ZNF385D, CXCL14, RGS12, SORCS3, KCNQ5, CNR1, and ADARB2. On the other hand, cluster I03 showed heterogeneous expression of those genes, while cluster I07 showed a similarly homogeneous expression pattern of those genes in some cells from the ADAD cases and all cells from the male RELN-COLBOS case included in that cluster (Fig. 3d).

**Figure 3.**
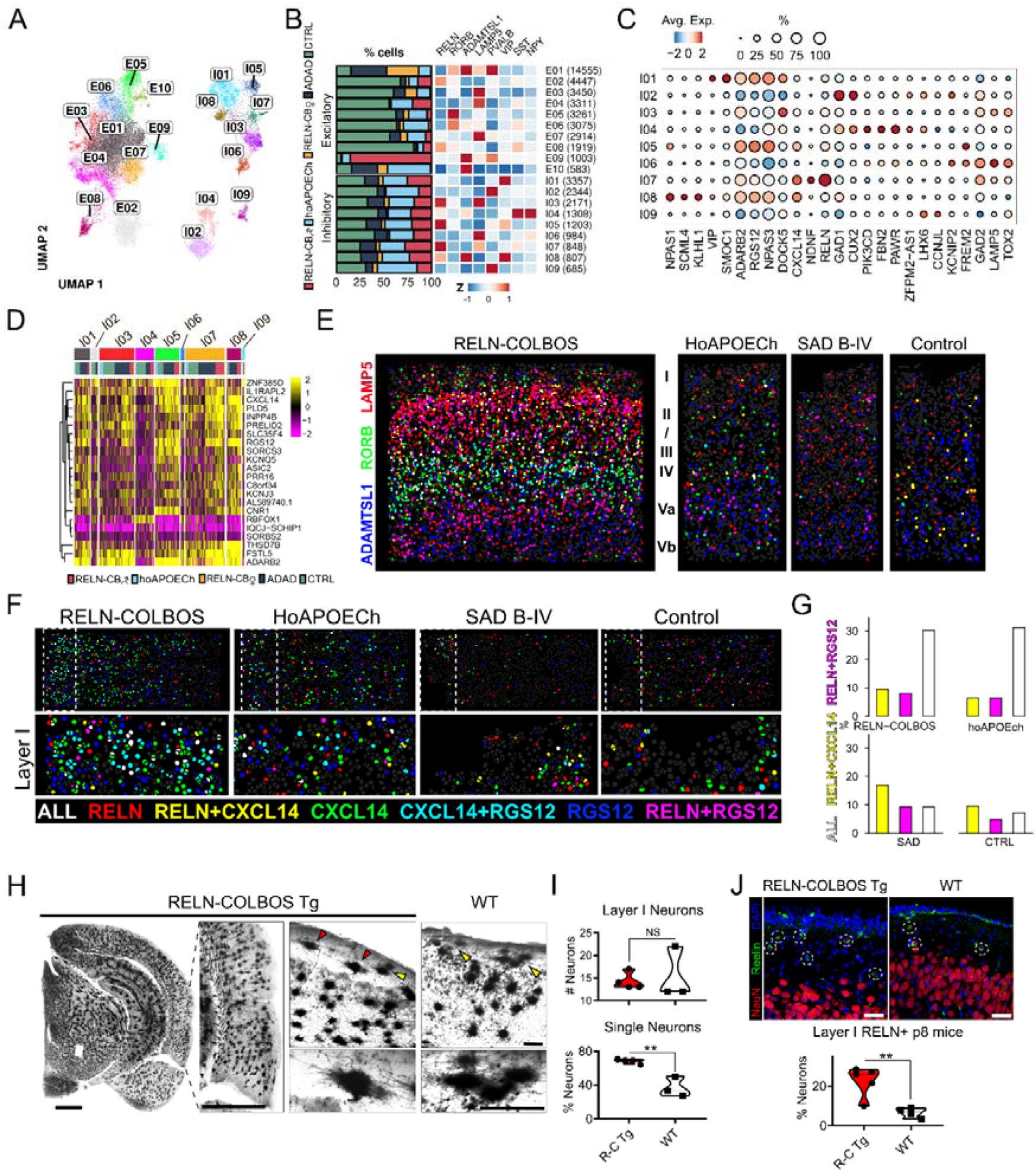
Characterization of unique RELN+ inhibitory Layer I neurons in the male RELN-COLBOS case. A. UMAP projection of single-nuclei transcriptome profiles from Excitatory (E) = 38518, and Inhibitory (I) = 13707 neurons. B. Stacked bar plot of neuronal subtype proportions, normalized by number of cases per condition, with a marker gene heatmap for excitatory and inhibitory neurons. C. Dot plot of canonical subtype marker expression and proportion in inhibitory neurons. D. Heatmap of top differentially expressed genes in RELN+ inhibitory neurons, comparing CXCL14+RGS12+ vs. CXCL14-RGS12- cells. E. Combinatorial single-molecule FISH (csmFISH) of the EC showing cells positive for layer markers; cells positive for 2+ markers are shown as blended primary marker colors. F. csmFISH of sections of the EC showing RELN, CXCL14, and RGS12; cells positive for 2+ markers are shown as blended primary marker colors. Layer I is magnified. G. Bar plot of the proportion of cells positive for each marker and colored as in (F), within Layer I. H. Representative microphotograp of Golgi-Cox staining of a coronal section from a RELN-COLBOS Tg mouse brain, with magnified view of the EC (bar = 500 μm). Further magnified images and comparison between RELN-COLBOS Tg and wild type (WT) Layer I bipolar neurons (bar = 50 μm). I. Violin plots for the quantification of the number of neurons detected in Layer I, and for the percentage of single bipolar neurons detected in this Layer in RELN-COLBOS Tg (RC-Tg, n = 4) and WT (n = 3) entorhinal cortices stained with Golgi-Cox, ** = p< 0.01. J. Representative immunofluorescent images of RELN-COLBOS Tg and WT Layer I of EC from p8 mice, stained for NeuN (red), Reelin (green), and DAPI (blue), bar = 50 μm. Below is the violin plot for the percentage of Reelin positive neurons detected in Layer I (outlined in white dotted lines) in RC-Tg (n = 4) and WT (n = 5), ** = p< 0.01, t-test.

To better ascertain the specific location, and thus the possible role of RELN+ neurons identified in the ADAD protected cases, we performed another round of spatial transcriptomics. At this instance we chose combinatorial single molecule Fluorescence In Situ Hybridization (csmFISH) for a panel of 64 genes, including genes either known to be cortical Layer markers, or as cluster identity markers in our snRNA seq experiments (Suppl. Table 5). Given the variability of postmortem intervals and formalin fixation times among our samples, just a subset of cases could be used and showed significant signals. This subset included the male RELN-COLBOS case, the hoAPOECh case, and three control cases with various degrees of AD pathology severity. Markers such as LAMP5, RORB, and ADAMTSL1 allowed for the identification of the cortical Layers in these samples. The male RELN-COLBOS case showed a higher density of transcripts, and more distinctive signal distribution among layers than the other subjects (Fig. 3e, Suppl. Fig. 4). Consequently, we used csmFISH to define the location of the RELN+ cells we identified in the snRNAseq analysis. We used RELN, RGS12, and CXCL14 as markers for inhibitory neurons of interest. As expected, cells simultaneously expressing these three genes were predominantly localized in Layer I, and in ADAD protected cases (Fig. 3f-g).

For further clarification of the impact of the RELN-COLBOS mutation in Layer I neuronal organization, we performed Golgi-Cox staining in 6 months old transgenic RELN-COLBOS mice (RELN-COLBOS Tg) ^1^ and littermate wild type controls. Interestingly, we identified differences in the distribution of Layer I neurons between transgenic and wild type mice (Fig. 3h). Bipolar interneurons in Layer I showed to be significantly more isolated in RELN-COLBOS Tg mice, while they presented in clusters in their wild type counterparts (Fig. 3h-i). The molecular signature of RELN+ cells identified in the male RELN-COLBOS case using snRNAseq and csmFISH, together with the neuronal morphology and distribution pattern in 6 months old RELN-COLBOS Tg mice suggest that the RELN-COLBOS mutation might influence RELN+ cells functionality and development. Thus, we evaluated Layer I RELN+ neurons density in the EC of p8 RELN-COLBOS Tg and wild type mice. We identified a significant increase in RELN+ neurons proportion of Layer I cells in the RELN+ COLBOS Tg (Fig. 3j), suggesting that cellular changes induced by the RELN-COLBOS mutations in inhibitory cells can occur already during neurodevelopment since they can be detected so shortly after birth.

### Specific excitatory neuronal subtype in lower cortical Layers in the male RELN-COLBOS patient

RELN+ neurons were also identified among excitatory cells, in clusters E02, E07, E08, and E09 (Fig. 4a). Remarkably, excitatory cluster E09 was almost exclusively composed by neurons from the male RELN-COLBOS case (Fig. 3b). These neurons also showed expression of SNTG2, ADAMTSL1, LRP4, LRP6, DAB1, KCNB2, FGF13, ATP7B, CUX2, and CLSTN2. Out of these markers, SNTG2 and ADAMTSL1 showed a more specific expression pattern for this cluster (Fig. 4a), suggesting their identity as pyramidal neurons in Layers II/II and Layer V (Allen Brain Atlas) ^43^. Additionally, E09 neurons from the male RELN-COLBOS case show a distinctive expression signature in this cluster, including ARL17B, STNG2, and LINC02607 (Fig. 4b), further supporting their possible location in Layer V. In fact, ADAMTSL1, ARL17B, and SNTG2 positive neurons were shown to belong mostly to cluster E09 and to the male RELN-COLBOS case (Fig. 4c). To further elucidate the identity of these neurons, we used the markers of cluster E09 that would show Layer specific distribution in our csmFISH panel. Thus, we used DAB1, mostly distributed in superior Layers, CLSTN2, mostly distributed in lower Layers, and LRP6, strongly expressed in this cluster (Fig. 4d, Suppl. Fig. 4). In accordance with the snRNAseq results, the male RELN-COLBOS case showed increased numbers of neurons expressing these three markers, either simultaneously or individually, distributed in Layers II/III to Layer Va (Fig. 4e).

**Figure 4.**
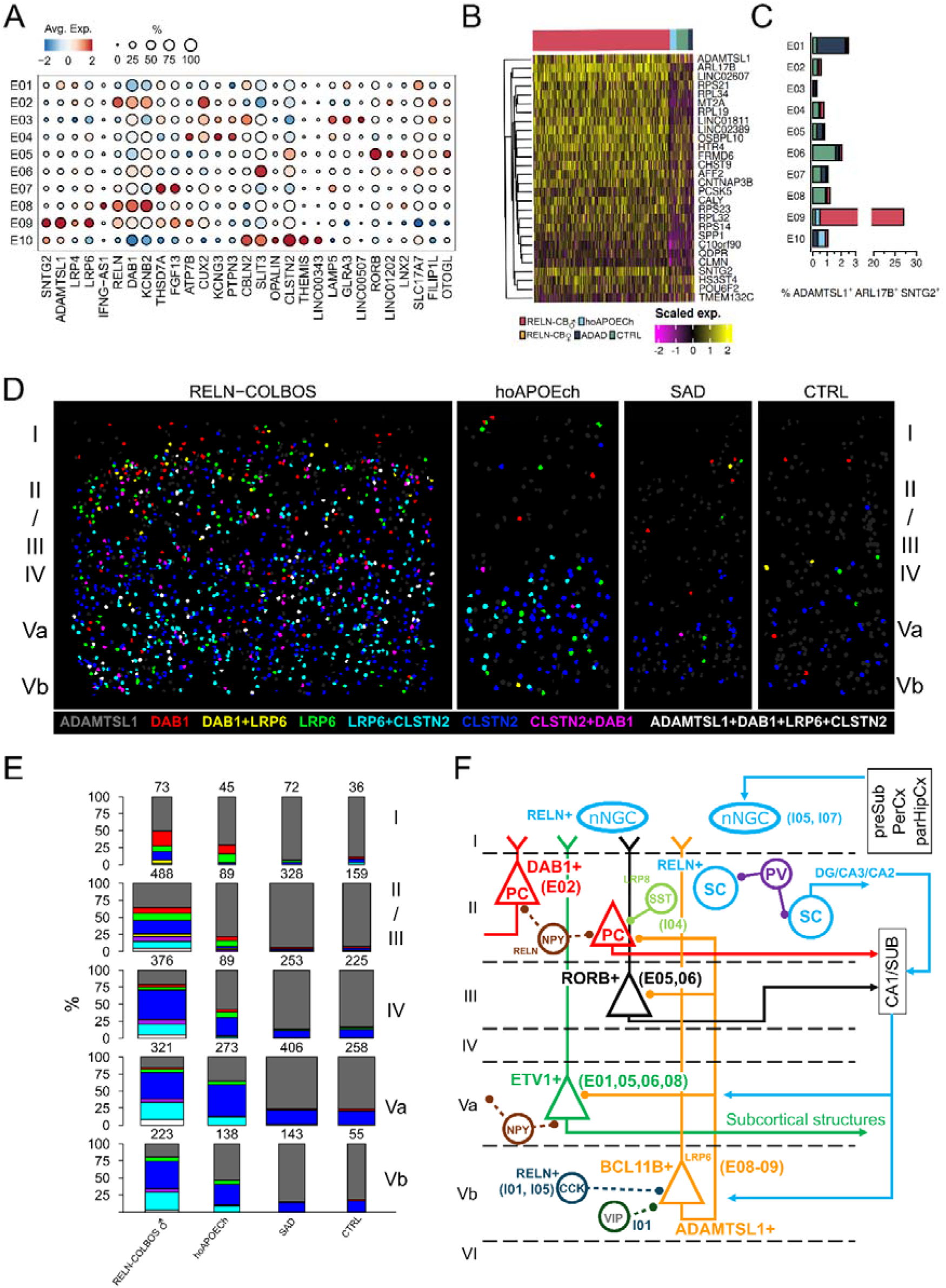
Enrichment of ADAMTSL1 positive cells in the male RELN-COLBOS case. A. Dot plot of canonical subtype marker expression and proportion in excitatory neurons. B. Heatmap of top marker gene expression for the E09 cluster. C. Bar plot of cell proportions positive for distinctive E09 markers (ADAMTSL1, ARL17B, SNTG2). D. csmFISH of the EC showing DAB1, LRP6, and CLSTN2 in ADAMTSL1+ cells only; cells positive for 2+ markers are shown as blended primary marker colors. E. Stacked bar plots of cell proportions positive for markers and colored as in (D), by cortical layer; bar width reflects the total number of ADAMTLS1+ cells per layer. F. Scheme of neuronal circuitry organization between entorhinal cortical layers. Different neuronal types are colored and denominated by their marker genes. The snRNA seq neuronal clusters in which they were distributed are written within parentheses.

Our analysis indicates that secreted levels of Reelin in the male RELN-COLBOS case had a strong localized protective effect in the EC of the male carrier of this mutation. This localized effect is reflected by increased abundance of RELN+ non-neurogliaform inhibitory neurons in Layer I, and RELN+ excitatory pyramidal neurons in Layers II/III and Layer Va. These changes likely influenced the functionality of the EC circuitry and its signaling output to other hippocampal regions and other associated cortical and subcortical structures, while affecting less its intrinsic EC networks ^44,45^ (Fig. 4f), conferring resilience to AD pathological changes.

### RELN-COLBOS mutation localized effect in glial cells via non-canonical LRP receptors

Next, we examined the expression pattern of astrocyte and oligodendrocyte subclusters to identify any possible difference in the RELN-COLBOS carriers. We identified seven astrocytic clusters (Fig. 5a), seemly with three expression patterns. Astrocyte clusters Ast01 and Ast05 showed similar groups contribution and high expression of CADM2, CACNG2, KCNH1, KCNH7, KCNJ6, GFAP, RELN, and HSPA8. In addition, cluster Ast01 also showed expression of HSP90AA1, and HSP90AB1, similar to the PSEN1 E280A ADAD astrocytic signature previously reported in frontal cortex ^46^. Clusters Ast03 and Ast06 showed similar expressions of NRXN1, PTN, AQP4, SLC1A2, GPC5, GLUL, and SLC1A3. However, Ast06 also showed expression of HSP90AA1, EEF1A1, UBB, UBC, RAC1, RAB11A, TUBB4A, and strong expression of HSP90AB1. Finally, Ast04 showed expression of UBB, HSPB1, S100B, TNC, CD44, PTN, AQP4, GLUL, SLC1A3, and strong expression of C3. Therefore, clusters Ast01, Ast04, and Ast05 showed a more reactive phenotype, particularly Ast04. Meanwhile, cluster Ast03 showed a more homeostatic phenotype, with cluster Ast06 showing an intermediate phenotype between both states (Fig. 5b). We performed pseudotime analysis to identify the change between homeostasis and reactivity in the astrocyte clusters. Cluster Ast03 was defined as the homeostatic origin while cluster Ast04 shown the furthest distance from it, in alignment with its strong reactive profile (Fig. 5c). Interestingly, cluster Ast03 shows a relatively high contribution of male RELN-COLBOS cells (Fig. 5b), suggesting that EC protection is associated with a homeostatic astrocyte phenotype in this case.

**Figure 5.**
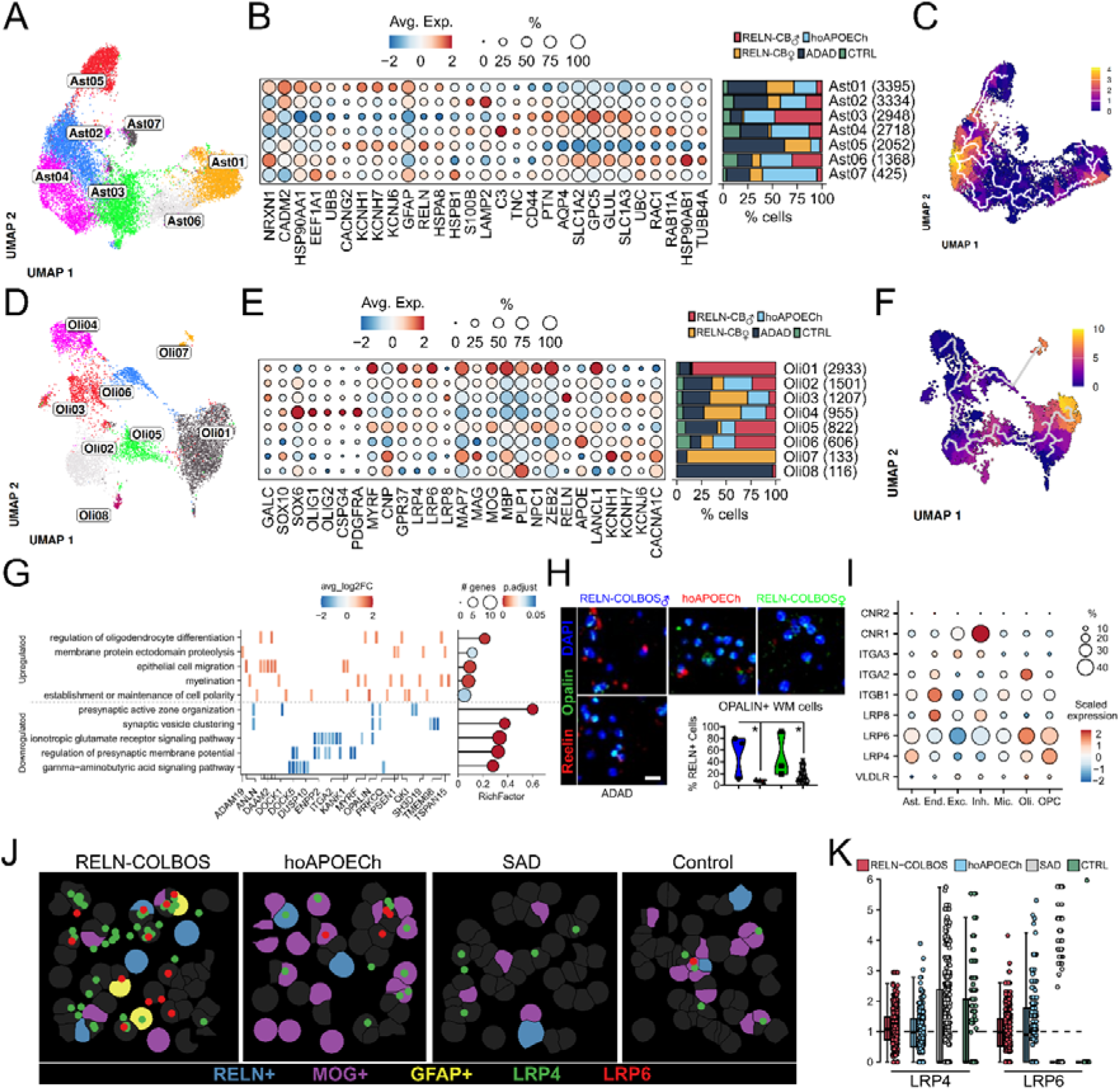
Localized effect of mutated Reelin in glial cells via LRP6 receptor in ADAD protected cases. A. Uniform manifold approximation and projection (UMAP) projection of snRNA-seq data from astrocytes. B. Stacked bar plot of proportions of astrocytic clusters with astrocytes markers dot plot. C. UMAP visualization of the astrocytes trajectory colored by pseudotime computed in Monocle 3. D. UMAP projection of snRNA-seq data from oligodendrocytes. E. Stacked bar plot of proportions of astrocytic clusters with oligodendrocytes markers dot plot. F. UMAP visualization of the oligodendrocytes trajectory colored by pseudotime computed in Monocle 3. G. Top 5 biological processes from up and downregulated genes from the comparison between Oli02 and Oli01 clusters are shown as a heatmap for the gene expression and lollipop for the significance and term enrichment. H. Representative immunofluorescent images of hippocampal white matter from RELN-COLBOS cases, hoAPOECh case, and ADAD cases (n=5), stained for Reelin (red), Opalin (green), and DAPI (blue), bar = 20 μm. Below is the violin plot for the percentage of Reelin positive and Opalin positive cells detected in three representative areas at 20x magnification per case, * = p< 0.05, Annova test. I. Dot plot of scaled gene expression of canonical and non-canonical RELN receptors in the cell types. J. Magnification of molecular cartography showing the cells positive for RELN, MOG and GFAP and the transcripts of LRP4 and LRP6. K. Box plot of ratio of neighboring cells to RELN+ cells that contain LRP4 or LRP6 respective to the total number of cells that are positive to LRP4 or LRP6.

Then we analyzed oligodendrocytes and identified eight subclusters (Fig. 5d). Notoriously, cluster Oli01 and Oli07 were mostly composed by male and female RELN-COLBOS oligodendrocytes, respectively (Fig. 5e). Cluster Oli01 presented a distinctive expression signature with MYRF, GPR37, LRP4, LRP6, MAP7, MAG, MOG, MBP, PLP1, NPC1, ZEB2, and LANCL1. This expression profile was similar to that from another cluster with a relatively high contribution of male RELN-COLBOS cells, Oli05, with the additional expression of CNP, and APOE. Cluster Oli07, mostly composed by female RELN-COLBOS oligodendrocytes, showed relatively high expression of CNP, MAP7, MAG, MBP, ZEB2, KCNH1, KCNH7, KCNJ6, and CACNA1C. Finally, cluster Oli04 showed high expression of SOX10, SOX6, OLIG1, OLIG2, CSPG4, and PDGFRA (Fig. 5e). These markers identify this cluster as a maturing oligodendrocyte, while clusters Olig01 show higher expression of mature / myelinated oligodendrocytes. In effect, pseudotime analysis shows cluster Oli04 as the origin, with cluster Oli01 as the more differentiated one (Fig. 5f). Differential gene expression analysis between clusters Oli01 and Oli02, with a relatively similar number of cells, showed enrichment of upregulated genes related to myelination function, while downregulated genes were associated with synaptic function (Fig. 5g, Suppl. Table 6). A protective function for the RELN-COLBOS mutation aligns with our previous findings of RELN positive signal in white matter cells and more dense white matter in the male RELN-COLBOS carrier ^1^.

Intriguingly, oligodendrocyte cluster Oli01 and excitatory neurons cluster E09, mainly composed by male RELN-COLBOS carrier cells, showed increased levels of LRP4 and LRP6 genes as markers (Fig. 5e). LRP4 and LRP6 belong to the same receptor family as the canonical receptors for Reelin, VLDLR and ApoER2 (also known as LRP8) ^47^. Thus, we examined the expression distribution of Reelin-associated receptors, either canonical, non-canonical, or associated ^48–50^. We saw no relevant expression of VLDLR in any cell type, while LRP8 was expressed only in endothelial cells and inhibitory neurons. The latter showed also strong expression of cannabinoid receptor CNR1. Astrocytes showed relatively high expression of LRP4 and LRP6 receptors, and integrin receptor subunit ITGB1. Endothelial cells also showed high expression of integrin receptor subunits ITGB1, and ITGA2. Microglia cells only showed mild expression of ITGB1. Both oligodendrocytes and OPC showed strong expression of LRP4 and LRP6, while oligodendrocytes show strong expression of ITGA2 as well (Fig. 5i). As we could see in excitatory neurons and oligodendrocyte subcluster gene expression, relatively high LRP4 and LRP6 expression is associated with male RELN-COLBOS cell contribution. Thus, we used csmFISH data to identify the association between RELN+ cells and the distribution and identity of cells expressing LRP4 and LRP6. In effect, astrocytes and oligodendrocytes showed expression of both receptors in cells surrounding RELN+ cells in the male RELN-COLBOS case, and oligodendrocytes in the hoAPOECh case (Fig. 5j., Suppl. Fig. 5). The quantification of the proximity and abundance of LRP4 and LRP6 positive cells surrounding RELN+ cells showed that LRP4 expression surrounding RELN+ cells was universal, while LRP6 expression was restricted to ADAD protected cases (Fig.5k), suggesting a non-canonical binding between Reelin and LRP6.

### Oligogenic RELN-COLBOS associated protective effects via its non-canonical receptors

The possibility that the RELN-COLBOS mutation alters binding to non-canonical receptors may explain its observed effect in oligodendrocytes and white matter ^1^. Therefore, we tested this hypothesis by modelling the interaction between Reelin and LRP6 using AlphaFold3 and the protein-protein interaction algorithm (PrePPI) to align and compare the predicted Reelin-Lrp6 and Reelin-Lrp4 interactions to the known Reelin-Lrp8 interacting complex (Suppl. Fig. 6). Cavity analysis further predicted that the C terminal region of Reelin shares a protein interaction pocket with LRP6 (Fig. 6a). To validate that Reelin binds to Lrp6 receptor, we used synthetic Reelin C terminal region (CTR, aa 3427 to aa 3460), with or without the RELN-COLBOS mutation (H3447R) to recombinant Lrp6 receptor. As a positive control, we quantified, Dickkopf-related protein 1 (DKK1) binding to Lrp6 receptor, being DKK1 a known Lrp6 ligand ^51^. Wild type Reelin CTR shows significantly less binding affinity to LRP6 compared to DKK1 binding affinity. Conversely, RELN-COLBOS mutated Reelin CTR showed a similar binding affinity to that of DKK1 (Fig. 6b-c).

**Figure 6.**
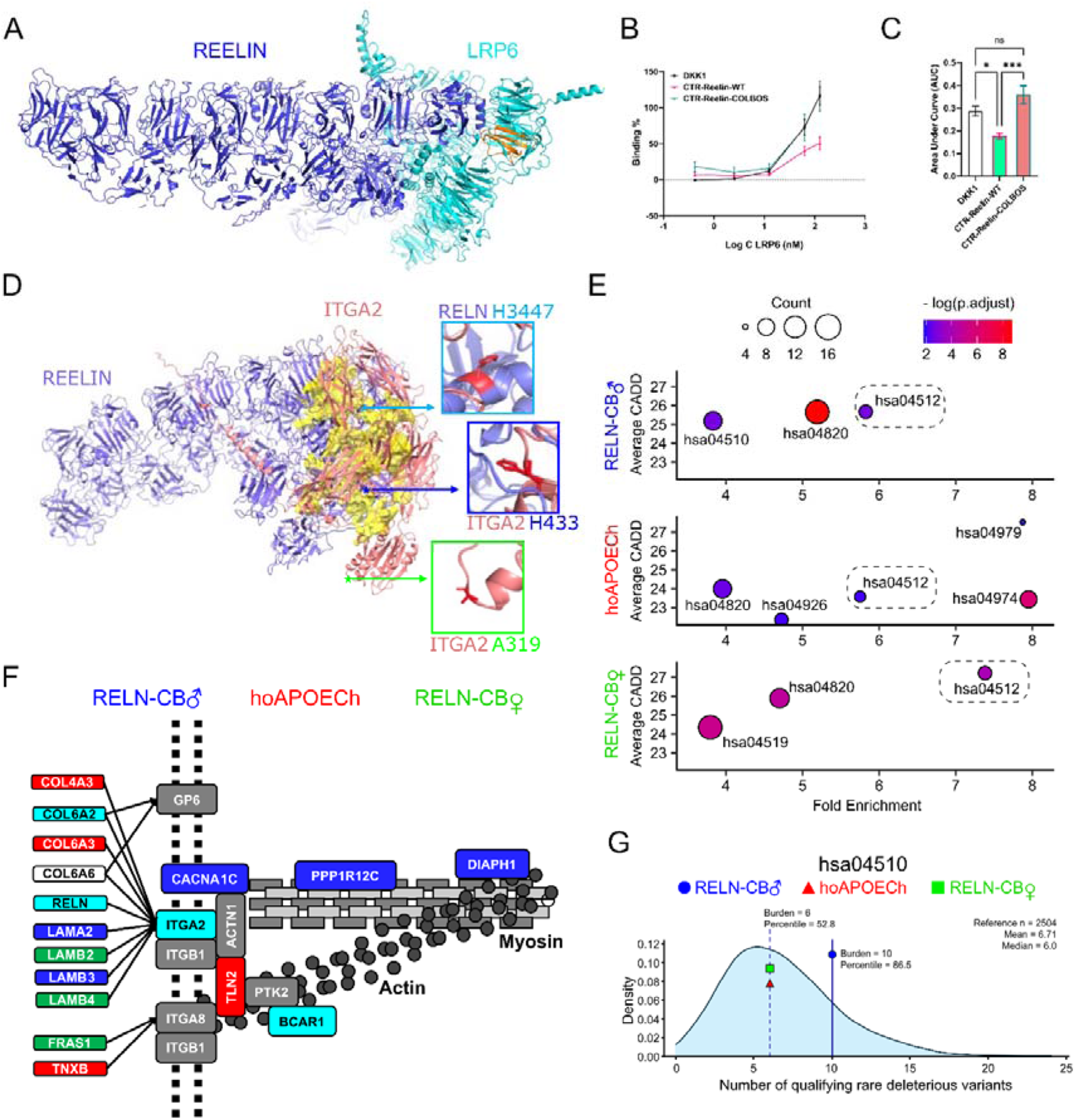
Non-canonical ligand-receptor interactions for Reelin and oligogenic mutational effects in ADAD protected cases. A. PrePPI predicted RELN and LRP6 aligned with Alphafold3 predicted complex. Based on the crystal structure of the RELN–LRP8 complex (PDB: 5B4X), PrePPI predicts interactions between RELN and LRP6 (blue/orange), and between RELN and LRP4. These predicted interactions are further supported by AlphaFold3 (blue/cyan). In the LRP8 complex, RELN binds to the EGF-like and LDL receptor A domains - structural motifs also conserved in LRP4 and LRP6 (Suppl. Fig. 6). B. ELISA dose-response curves showing Dkk1, CTR-Reelin-WT, and CTR-Reelin-COLBOS binding to LRP6 across increasing concentrations. C. Quantification of the area under the curve (AUC) reveals significantly higher CTR-Reelin-COLBOS binding to LRP6 compared to CTR-Reelin-WT. Data represent mean ± SEM; one-way ANOVA with Tukey’s post hoc test; * (p < 0.05), ** (p < 0.001); ns = not significant. D. Alphafold3 predicted RELN (blue) and ITGA2 (pink) complex, with separately calculated cavities in ITGA2 highlighted in yellow. The locations of three mutations of interest are shown in the enlarged inset panels. E. KEGG pathway enrichment for rare exonic variants found in RELN-COLBOS♂, RELN-COLBOS♀, and hoAPOECh cases, plotted by enrichment ratio and average CADD (Combined Annotation Dependent Depletion) PHRED score. The ECM-receptor interaction pathway (hsa04512) is highlighted across all three groups. F. Pathway diagram of hsa04150 / hsa04512 pathways mapping mutated genes to the corresponding cases. The genes that are mutated in 2 or more cases are shown as the combination of primary marker colors. G. Rare exonic deleterious variants burden density plot for 1000g whole genome sequences obtained from 2504 subjects for the hsa04510 pathway. The RELN-COLBOS carriers (male = blue symbol, female = green symbol) and the hoAPOECh (red symbol) carrier burden are depicted relative to general population density.

While the first report of these cases suggested that the protection was monogenic ^1,52^, a deeper genetic analysis revealed that both cases also carried other exonic rare variants. Therefore, we tested whether any of these variants could contribute to the resilience phenotype observed in these individuals. The male RELN-COLBOS case also carries 909 exonic mutations with a global allelic frequency of 0.01 or less, the hoAPOECh case also carries 819 exonic mutations with a global allelic frequency of 0.01 or less, and the female RELN-COLBOS case also carries 656 exonic mutations with a global allelic frequency of 0.01 or less (Suppl. Table 7). We inspected these gene variants and found that both RELN-COLBOS carriers had a different mutation each in the non-canonical Reelin receptor ITGA2, a possible candidate receptor for Reelin and highly expressed in oligodendrocytes in our dataset (Fig. 5i). The male RELN-COLBOS patient was heterozygous for mutation H433Q in ITGA2, while his sister was heterozygous for mutation A319T in ITGA2. Similarly to LRP6, AlphaFold3 modelling of Reelin-ITGA2 showed that the C terminal region of Reelin shares a protein interaction pocket with ITGA2. Interestingly, only mutation H433Q in ITGA2 is also located in the protein interaction pocket (Fig. 6d).

Our data strongly suggests that extreme protection against AD is a result of multiple rare structure-modifying mutations impacting a common pathway and that the size of phenotypic effect of a gene variant can be potentiated if interacts directly with another mutated protein. The male RELN-COLBOS carrier can be an example of this multigenic-driven resilience against ADAD. However, not all exonic variants have the same deleterious effect. To determine the possible impact of these mutations we used CADD which is a tool developed for scoring the likelihood that single nucleotide variants, multi-nucleotide substitutions, as well as insertion/deletions variants in the human genome may alter the natural function of the encoded protein ^27^. The RELN-COLBOS mutation has a CADD score of 25.5, the APOECh mutation has a CADD score of 25.3, the ITGA2 H433Q mutation has a CADD score of 22.8, and the ITGA2 A319T mutation has a CADD score of 17.06. Once filtered for a CADD score equal or higher than 20, the male RELN-COLBOS case carries 241 rare non-synonymous gene variants, the hoAPOECh case carries 209 rare non-synonymous gene variants, and the female RELN-COLBOS case carries 176 rare exonic gene variants (Suppl. Table 7, Suppl. Fig. 7). Gene enrichment analysis of these genes using the KEGG pathways database to detect direct functional protein interactions showed several enriched pathways enriched for each case. The ECM – receptor interaction pathway (hsa04512) was the top enriched pathway in both RELN-COLBOS carriers, simultaneously showing the highest average CADD score in the affected genes. Meanwhile, even though the Cholesterol metabolism pathway (hsa04979) was the one satisfying these characteristics in the hoAPOECh case, the hsa04512 pathway was also enriched with a high average CADD score. The ECM – receptor interaction pathway is complementary to the Focal Adhesion pathway (hsa04510), also enriched with high CADD average score in the male RELN-COLBOS carrier (Fig. 6e, Suppl. Fig. 8). Both pathways include extracellular matrix (ECM) proteins like collagens, laminins, and other proteins such as Reelin, as ligands for Integrin receptor heterodimers. Some of these genes also showed mutations with high potential changes in their corresponding encoded proteins (Suppl. Table 8). Downstream functions of this pathway include actin cytoskeleton related functions involved in cell motility, proliferation and survival ^53^. Both RELN-COLBOS carriers and the hoAPOECh carrier also carry highly deleterious exonic mutations in genes directly interacting in this biological pathway (Fig. 6f). To contextualize this finding, we analyzed rare exonic variants in the 1000g whole genome dataset by using the same filters (AF ≤ 0.01, CADD ≥ 20). Next, we studied rare exonic gene variants in the genes belonging to the hsa04510, hsa04512, and hsa04141 Kegg pathways, and calculated the individual burden for the 1000g sample. We used the distribution of gene variant burden in these pathways to assess the rare variant burden uniqueness of the two RELN-COLBOS carriers and the hoAPOECh carrier. The analysis showed that only the male RELN-COLBOS carrier diverged from the general population burden distribution for pathways hsa04510 (percentile 86.5) and hsa04512 (percentile 74.2) (Fig. 6g, and Suppl. Fig. 9), indicating another point of divergence between the two RELN-COLBOS carriers. These pathways are among the most common to include rare exonic variants in the general population (Suppl. Table 9 and Suppl. Fig. 10). In summary, it is possible that RELN-COLBOS mutation protective effects are enhanced by other rare gene variants present in those carriers, and one possible mechanism affected by this oligogenic effect is the Integrin mediated focal adhesion pathway.

## DISCUSSION

Our results indicate that one single mutation in one gene can have a strong effect on different molecular phenotypes in different cell types. This effect can be further mediated by other genetic variants involved in molecular pathways that include the gene with the higher effect reported, ultimately resulting in a distinctive phenotypic effect in ADAD.

The original report for the RELN-COLBOS mutation focused on a male heterozygous carrier, also carrying the PSEN1 E280A mutation, presenting with delayed dementia onset of 25 years ^1^. However, even though the sister of this patient was also a heterozygous RELN-COLBOS carrier, her disease onset was delayed only by 9 years (1.5 standard deviations from the PSEN1 E280A population average), and she did not present either resistant or resilient phenotypes against AD pathology. For instance, she showed an EC neuronal density similar to that of other AD patients (Suppl. Fig 2). This phenotypic difference was originally attributed to sex-dependent differences in RELN expression ^1,54^. Our current results suggest that the male RELN-COLBOS patient was also a carrier for additional unique rare variants, not shared by his sister, and those resulted in either cis-gene effects via intronic mutations, or in oligogenic synergic effects in whole molecular pathways via additional exonic mutations. Our single nuclei and spatial transcriptomic results support this concept. The male RELN-COLBOS patient showed unique regional gene expression patterns in the EC, characterized by unique excitatory neuronal and oligodendrocyte populations, and specific gene expression profiles. Furthermore, our study shows a direct localized effect of the RELN-COLBOS mutation in neuronal and glial cells via non-canonical Reelin receptor pathways, some of them also affected by additional mutations. This unique genotype-phenotype association was not evident in the female RELN-COLBOS patient, whose additional rare genetic variants did not potentiate the RELN-COLBOS protective effect as compared to her brother.

Bulk spatial transcriptomic analysis of several hippocampal regions showed a common gene expression profile for both the male RELN-COLBOS case and the homozygous APOECh case. This profile was not shared by the EC of the male RELN-COLBOS case. Nevertheless, this commonality shared by the two protected cases could indicate a common molecular pathway of protection, for instance via Dab1 signaling, as previously suggested ^1^. On the other hand, besides the protection evidenced by the delayed dementia age of onset, these cases shared another feature, a short duration of disease of two years before death by AD unrelated causes. Hence, it is also possible that the common gene expression pattern found in the hippocampal regions of these two cases reflects short disease duration instead. This possibility cannot be discarded, given that such disease duration is per se uncommon in the PSEN1 E280A population, and that no donated PSEN1 E280A brains shared that characteristic at the time of the study. In either case, the gene expression pattern detected, including upregulation of synaptic function, signaling, protein catabolism, and metabolism, aligns with early stages of AD pathology ^40^, suggesting activation of compensatory mechanisms against neurodegeneration ^55–58^.

The transcriptomic profile of the EC, using snRNAseq and GAEC analysis, revealed a high number of eGenes associated with rare genetic variants carried by the male RELN-COLBOS case. These downregulated genes included calcium and potassium channels, and GABA receptors, together with other genes involved in synaptic and signaling cascades. Interestingly, the expression of these eGenes associated mostly in glial cells. Subsequent analysis indicate that some of these genes are highly expressed in astrocyte and oligodendrocyte clusters that show evidence of pathological processes such as astrocytic over-reactivity ^46^, or evidence of myelination dysfunction, as exemplified by upregulation of KCNJ6 ^59^. One caveat of this analysis is not only our low sample number, but the intrinsic limitations associated with the study of rare and ultra rare genetic variants ^60^. So, even though we restricted our GAEC analysis to cis-gene associations, it is not possible to identify associations for unique single nucleotide variations within a gene, because by being carried by a single individual all show the same degree of association. However, given that the association between gene variants and expression was not uniformly detected in all the cells studied for each individual case, and that they could show association for one or more eGenes in different cell types in the same individual, eGenes detected can be considered as effective eGenes for all cis-gene variants identified in each gene. We deem these findings to be hypothesis generating. Therefore, the possible role of these genetic variants on astrocytic or oligodendrocytic phenotypes in Alzheimer’s pathology should be further evaluated in a larger sample of cases, including more than one individual carrying them.

On the other hand, even though neurons showed a low number of eGenes, the analysis of neuronal clusters revealed distinctive neuronal populations specific for the male RELN-COLBOS case, further confirmed by spatial transcriptomic analysis. This case showed a high number of RELN+, CXCL14+, RGS12+ inhibitory neurons in Ec cortex Layer I, when compared to the hoAPOECh case, other ADAD cases and healthy controls. Their transcriptomic profile and location suggest they are non-neurogliaform cells, previously characterized in adult mice ^42^. We confirmed anatomical changes in Layer I neuronal population in RELN-COLBOS Tg mice, showing higher proportion of single bipolar interneurons, opposite to the most common presentation of cells in clusters in wild type mice. More importantly, the male RELN-COLBOS case showed almost exclusively a subtype of excitatory neurons characterized by expression of ADAMTSL1, ARL17B, and SNTG2. Additionally, by using DAB1 and LRP6 as excitatory neuron cluster and cortical Layer markers, ADAMTSL1 positive neurons were mainly identified in Layers II/III and Va. An increased number of RELN+ cells in Layer I and increased number of excitatory ADAMTSL1+ pyramidal neurons in the male RELN-COLBOS case suggest a reinforcement of the inter Layer entorhinal circuitry, or reflects alterations in the modulation of main inputs into Layer I. This idea is further supported by high expression of LRP6 in the ADAMTSL1+ pyramidal neurons, that together with our finding of increased affinity binding between RELN-COLBOS mutated Reelin and LRP6 as non-canonical receptor, suggests a localized effect of the RELN-COLBOS mutation in EC Layer Va. This LRP6-dependent effect can also be seen in a subpopulation of fully differentiated oligodendrocytes, almost exclusively composed by cells belonging to the male RELN-COLBOS case and, showing high expression of LRP6 and myelination-associated genes. Further support for a localized effect of both RELN-COLBOS and APOECh mutations in the EC was the increased number of LRP4 and LRP6 positive cells surrounding RELN+ cells, with the additional effect of the RELN-COLBOS mutation in Reelin binding affinity to LRP6.

Our findings up to this point do not fully explain the differences between the male and the female heterozygous RELN-COLBOS carriers. The female case did not show any of the outstanding findings in either neuronal or glial cells. There was only one subpopulation of oligodendrocytes that was mainly composed by cells from the female RELN-COLBOS case, and it did not show the differentiated mature profile evidenced in the male case. Our GAEC analysis showed that, even though both RELN-COLBOS cases carried additional rare gene variants, the majority of eGenes associating with gene expression changes belonged to the male RELN-COLBOS case. In addition, we previously reported evidence of preserved white matter integrity in this case ^1^, further supported by our current findings regarding its differentiated oligodendrocytic profile. Intriguingly, the male RELN-COLBOS case also showed high expression of ITGA2 receptor in oligodendrocytes and carried a non-synonymous mutation potentially affecting its binding to its ligands. Both findings, mutations associated with downregulated eGenes, and the ITGA2 mutation, suggest that white matter integrity in the male RELN-COLBOS case was not only due to the RELN-COLBOS mutation, but to the aggregated effect of additional mutations in other genes involved in myelogenesis ^61,62^. Thus, although it is likely that the RELN-COLBOS mutation conferred some degree of protection to the female RELN-COLBOS carrier, as her age of onset was nine years over the PSEN1 E280A population average value ^63^, it was not enough to produce the outstanding effect observed in her brother, who also carried mutations in other key genes influencing oligodendrocyte differentiation and myelination.

These findings suggest an oligogenic protective effect on the RELN-COLBOS male case. In fact, the possible impact of simultaneous nonsynonymous mutations in ligand-receptor interactions suggest their additive effect in different molecular and cellular pathways. A second look at enriched molecular pathways affected by highly deleterious exonic mutations in the RELN-COLBOS cases and the hoAPOECh case showed PI3K-AKT, Focal adhesion, and Integrin signaling pathways commonly affected in the three cases, with the Integrin receptors signaling pathway as the most affected in both RELN-COLBOS carriers. Interestingly, this pathway was not the most enriched in the hoAPOECh case, but rather the cholesterol metabolism pathway, with additional mutations in NPC1, LIPG, and ANGPTL4. All three cases showed mutations in genes belonging to the Integrin signaling / Focal adhesion pathway, including integrins ligand COL6A6, mutated in all three cases with different variants. Integrin signaling is a main regulator of oligodendrocyte maturation, survival and myelination, and it is also a mediator between ECM signals and cytoskeleton remodeling ^62^. Thus, deleterious mutations in this pathway could affect oligodendrocytic response to other ongoing neurodegenerative processes in ADAD and potentiate or interfere with protective effects of other mutations. These findings in the ADAD protected cases were further contextualized when contrasted with rare exonic variants detection and KEGG pathways enrichment analysis in the 1000g dataset. Both Integrin-related pathways hsa04512 and hsa04510 present enrichment for rare exonic deleterious variants in general population. Nevertheless, the RELN-COLBOS male case presented a higher number of these variants relative to general population, suggesting that even among uncommon variants, the potential functional impact of the mutations involved might define the oligogenic effect in the pathway. A deeper analysis involving the relationship distance between affected proteins in the pathway, together with individual structural analysis for each gene variant will allow the identification of the size of effect of the contributing variants within each pathway. This issue is exemplified by the presence of two different ITGA2 missense mutations in each RELN-COLBOS carrier, with only the ITGA2 H433Q mutation carried by the male RELN-COLBOS case potentially affecting the interaction with Reelin.

Even though the PSEN1 E280A population is the largest identified so far with a single causative mutation for ADAD, including more than 1300 carriers, it is not a large population from a genomic point of view, rendering some analysis either non-viable or unreliable. However, it is possible to identify extreme phenotypic outliers and propose genetic associations to single rare variants, if their possible biological effect is validated further ^1,52^. One factor to consider is the size of effect that can be attributed to a candidate mutation. For instance, even though some genetic variants have shown strong genetic association to AD risk due to their frequency in affected individuals, their size of effect is so small that they cannot be considered to be causative ^64^. On the other hand, the size of effect of the APOE4 haplotype is large enough to be considered as a risk factor for AD on its own ^65^. The validation studies for the APOECh mutation suggest a relatively strong modifying effect for it, similar to other APOE variants ^66–68^. In this study, we conducted deep phenotyping analysis in EC tissue of the RELN-COLBOS carriers, and we identified a strong molecular effect associated with Reelin signaling and neuronal populations benefiting from it. However, this effect might be restricted to the EC, as suggested by the lack of general protection against AD pathology, and might be difficult to model in other organisms or cell cultures, given its apparent reliance on EC circuit organization. Our findings suggest that oligodendrocyte homeostasis is an additional factor in the strong protection identified in the male RELN-COLBOS carrier and that it results from the added effect of additional rare mutations carried by this patient. This could explain why the protective effect of the RELN-COLBOS mutation could not be confirmed in other studies^69^. At this point, it is not possible to assign a size of effect to the additional mutations identified in this case given sample size, but further studies will clarify this issue and will allow us to eventually develop a polygenic or oligogenic risk score for presentation of disease modifiers in PSEN1 E280A ADAD. We propose an alternative design of such polygenic and oligogenic models, including only genes with direct molecular interaction in well-established molecular pathways.

Clear limitations of our study include sample size and the unavailability of additional carriers of the rare variants under study. Because some variants were present in only a single donor, the observed genotype-associated expression changes may partially reflect donor-specific effects rather than true variant-driven effects. Therefore, findings for ultra-rare variants should be interpreted as exploratory and require replication in additional carriers. In addition, ADAD protected cases are unique individuals with extremely rare genetic fingerprints, making it challenging to devise replication studies. However, the depth of our analysis can be considered as a personalized medicine approach, which at the end brings the advantage of a more comprehensive understanding of what makes any of those cases so unique.

In conclusion, genetic mutations acting as phenotypic modifiers of disease presentation in ADAD can be further enhanced by additional gene variants in protected individuals, contributing to their extreme phenotypes and opening new avenues of research for a better understanding of AD pathophysiology.

## Supporting information

Supplementary index and figures

## Acknowledgements

The authors would like to acknowledge the invaluable contribution of the Colombian families suffering from Familial Alzheimer’s disease for donating biological samples for this study. Also, to Dr. Joseph Arboleda-Velasquez at Harvard Medical School for RELN-COLBS transgenic mice brain samples, to Dr. Alberto Rábano at Fundación Cien, in Madrid Spain, for control human FFPE hippocampal tissue, and to Dr. Juliana Acosta-Uribe for the genomic sequence of the female RELN-COLBOS carrier. We thank the staff and technical support of the UKE imaging facility (UMIF). This work was funded by the Deutsche Forschungsgemeinschaft (DFG) with project number 458854216, and by the Wilhelm Emanuel Zach Foundation, which is administered by the Bürgerstiftung Hannover, both to D.S-F.

## Contributions

D.S-F designed the study, and together with Y.E-A designed data analysis strategies. Y.E-A performed transcriptomic data analysis. D.S-F, Y.E-A, A.R., C.A.V-G, and V.F. performed genomic data analysis. C.M, H.Z, and J.G-P performed protein structural modeling and analysis. H.Z, C.M, Z.S and R.T. performed variant pathogenicity analysis. D.S-F, A.K-F, and M.P.W analyzed neuronal populations and devised the EC circuit model. N.V-M, J.U, and M.A-T performed experiments. D.S-F, N.V-M, A-V, S.K, and M.G performed neuropathological and histological analyses. D.C-M and A-V collected and prepared human brain samples. D.A and A-V collected and analyzed clinical data from the patients. D.S-F, Y.E-A, A.K-F, M.P.W, V.F, H.Z, A.R, and C.M drafted the manuscript. All authors read and approved of the contents.

