## Supplementary index and figures for "Mechanisms of resilience to autosomal dominant Alzheimer’s disease via oligogenic modulation of rare variants in the entorhinal cortex"

Supplementary Material Index – El Amri et al. 2026

Supplementary Tables

| | |
|---|---|
| 1 | Demographic data of cases used in this study |
| 2 | List of enrichment terms and genes for bulk spatial transcriptomics DGE comparison analysis between protected and unprotected ADAD cases |
| 3 | List of genes and biological terms from the heatmap of common upregulated genes of hippocampal ROIs in ADAD protected cases (Figure 1h) |
| 4 | List of enrichment terms and genes from the chord diagram depicting functionally related eGenes identified using GAEC analysis of snRNAseq data, according to KEGG (A) and GO:BO (B) (Figure 2) |
| 5 | List of genes selected for the elaboration of the csmFISH panel (Resolve) |
| 6 | List of enrichment terms and genes derived from the comparison between Oligodendrocytic clusters 01 vs 02 (Figure 5g) |
| 7 | List of the number of rare exonic variants identified in ADAD protected and unprotected cases used in this study |
| 8 | Identification of potential deleterious effects of rare gene variants identified for the KEGG pathway hsa04510 in the RELN-COLBOS male carrier |
| 9 | Summarized results of KEGG pathways enrichment analysis in the 1000g dataset |

Supplementary Figures

| | |
|---|---|
| 1 | Density plots for transcriptomic profile of hippocampal ROIs (GeoMx) |
| 2 | EC neuronal density in RELN-COLBOS carriers, hoAPOECh carrier, ADAD cases and SAD cases |
| 3 | Manhattan plot of rare-variant-expression associations identified in neurons and other cell types |
| 4 | Full slide view of smFISH findings |
| 5 | Full slide view of LRP receptor transcripts localization relative to RELN+ cells via smFISH |
| 6 | PrePPI prediction of PPI of RELN/LRP6 vs RELN/LRP8 |
| 7 | Barplots rare exonic variants in protected and unprotected ADAD cases included in this study |
| 8 | Dot plots fore enrichment vs CADD scores of KEGG pathways for ADAD cases included in this study |
| 9 | Rare exonic deleterious variants burden density plot for hsa04512 and hsa04141 KEGG pathways |
| 10 | Bar plots for most common KEGG pathways presenting rare exonic variants in the 1000g dataset |

Suppl. Fig. 1

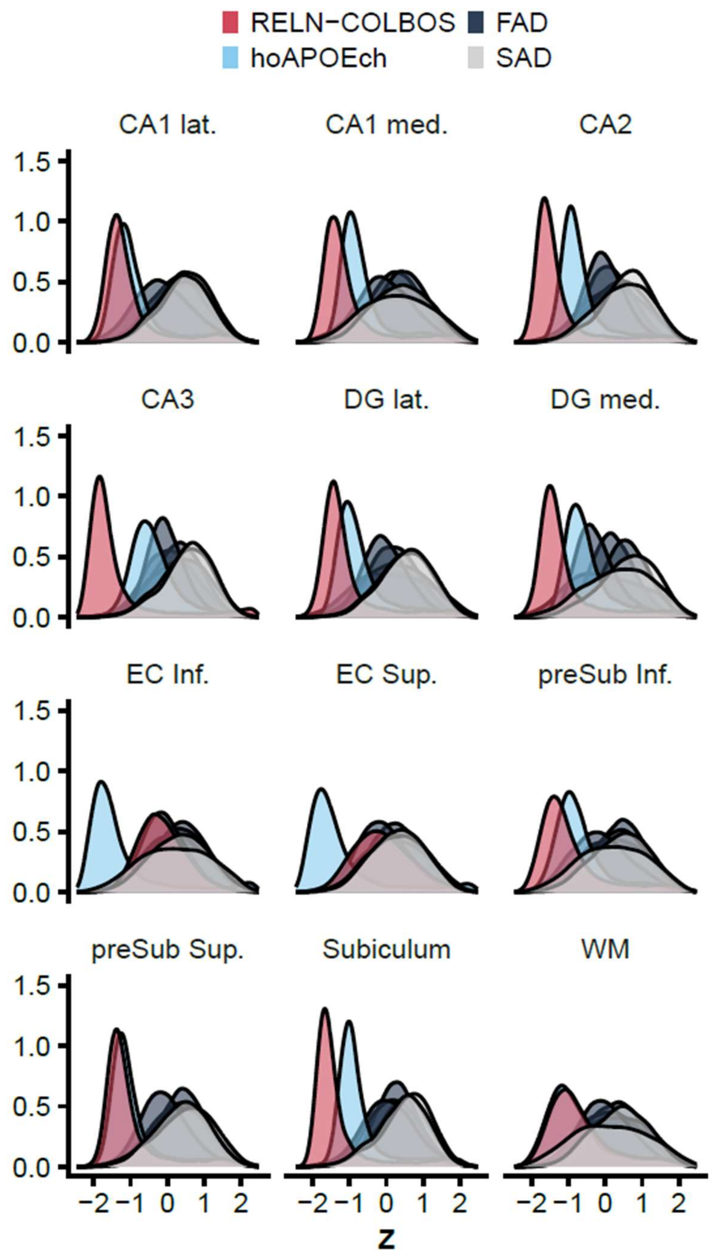


Density plots of individual gene expression across all genes in all hippocampal regions of interest. DG: dentate gyrus, EC: entorhinal cortex, preSub: presubiculum, WM: white matter, lat.: lateral, med.: medial, Inf.: inferior, sup.: superior.

Suppl. Fig. 2

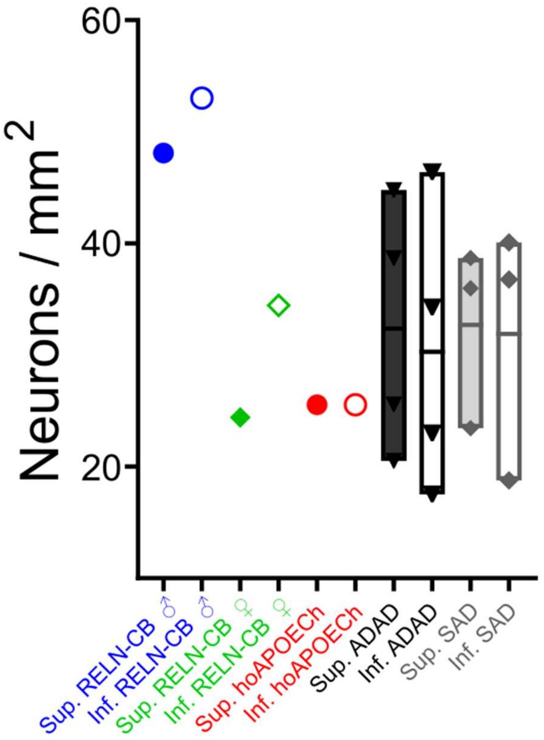


Bar graph for neuronal density measured in superior and inferior entorhinal cortical layers in RELN-COLBOS carriers, hoAPOECh carrier, ADAD cases (n=4) and SAD cases (n=3).

Suppl. Fig. 3

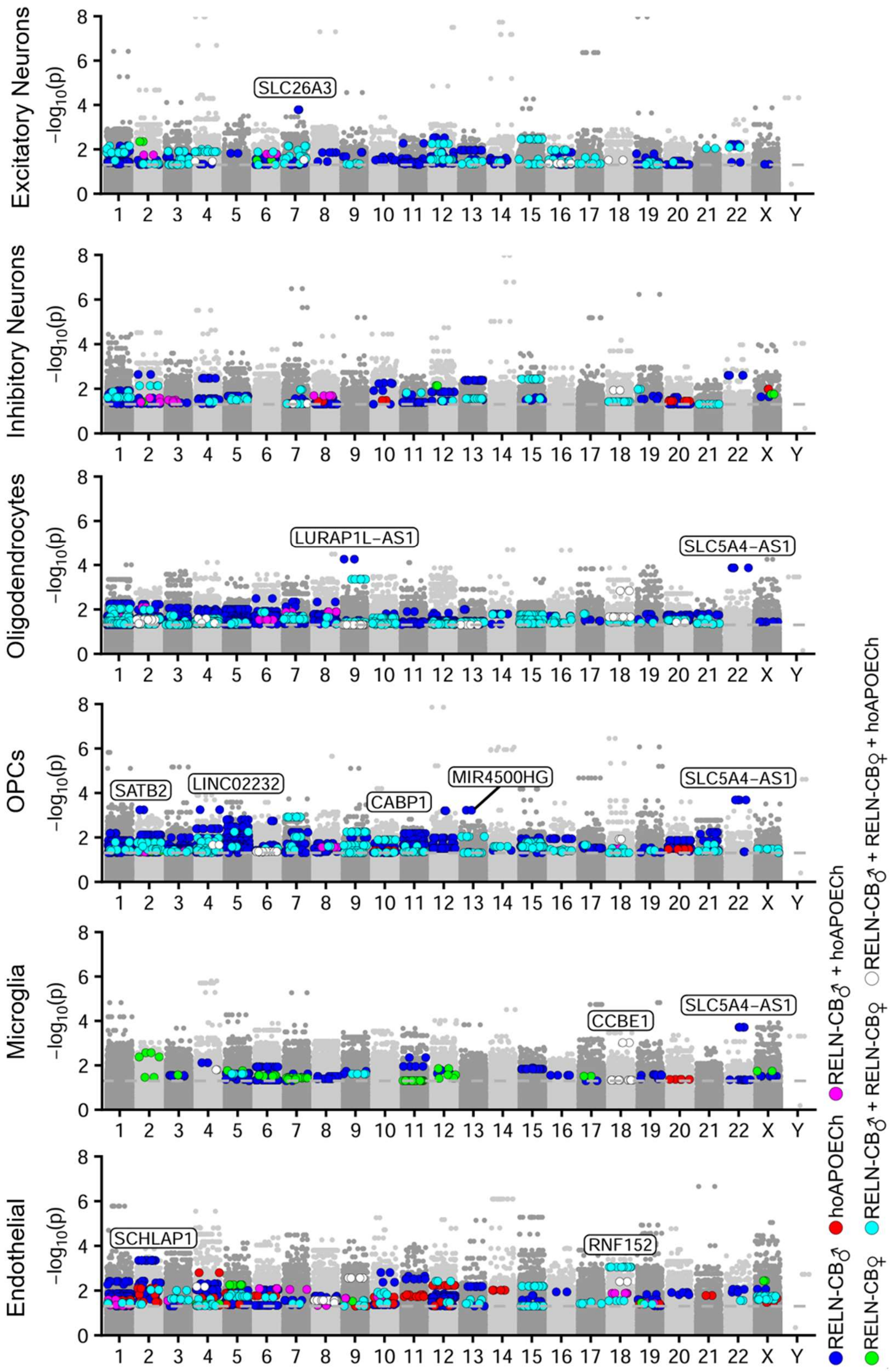


Manhattan plot of rare-variant-expression associations in snRNA seq cell type clusters.

Suppl. Fig. 4

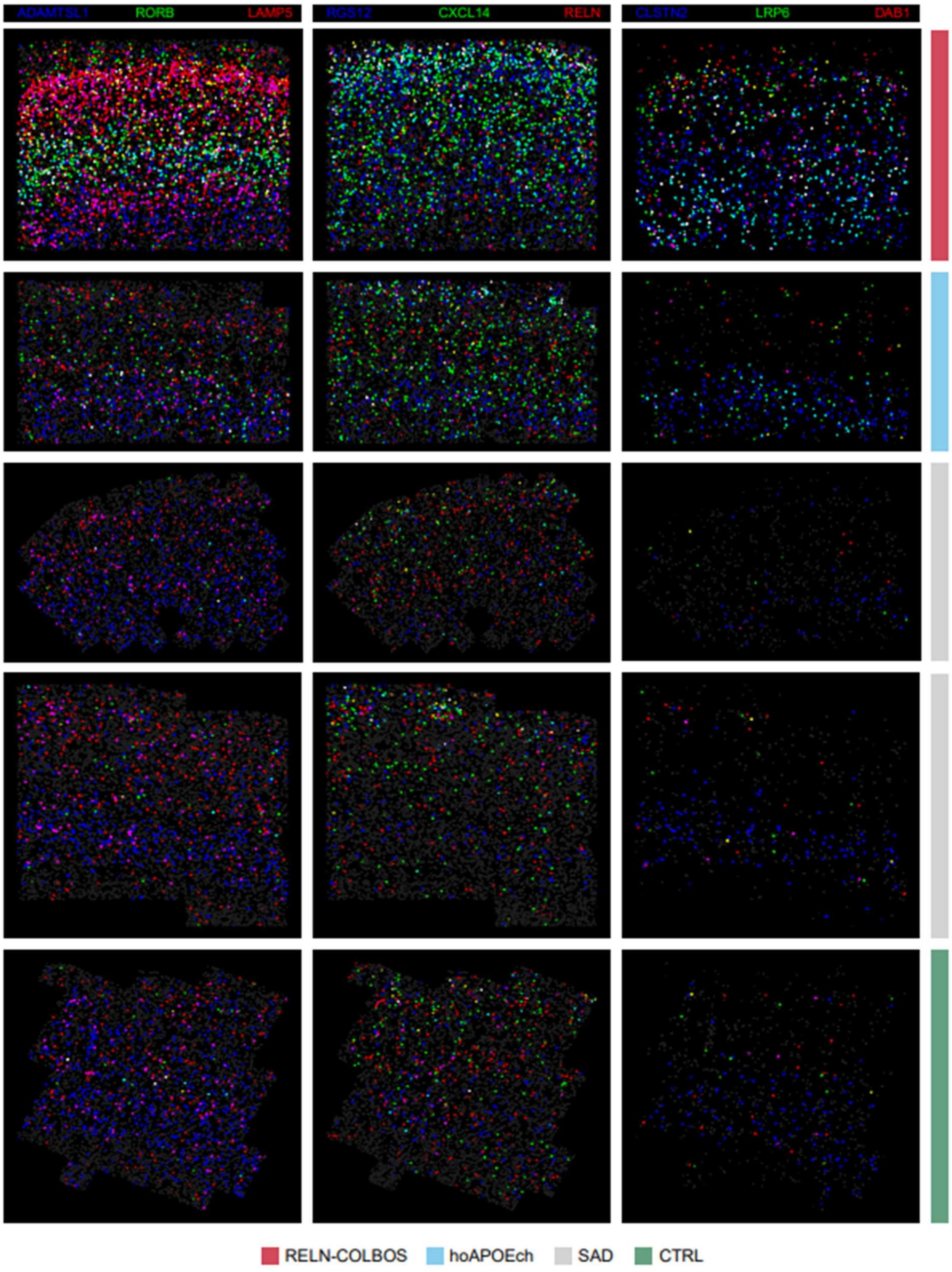


Combinatorial single-molecule FISH (csmFISH) of the EC showing cells positive for ADAMTSL1, RORB, and LAMP5 (left column), RGS12, CXCL14, and RELN (center column, and CLSTN2, LRP6, and DAB1 (right column). Cells positive for 2+ markers are shown as blended primary marker colours.

Suppl. Fig. 5

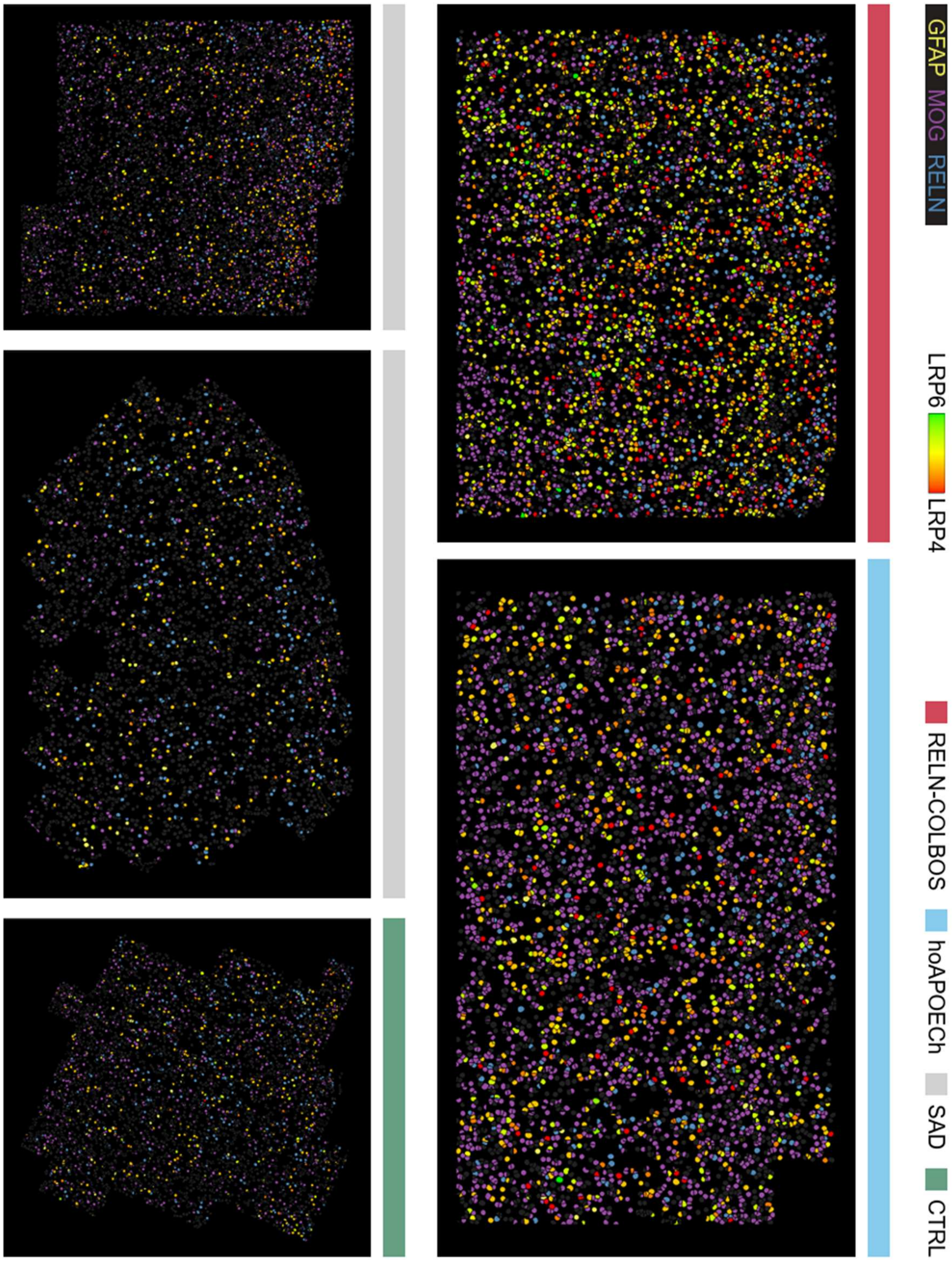


Combinatorial single-molecule FISH (csmFISH) of the EC showing cells positive for glial markers and LRP4 or LRP6 expression.

Suppl. Fig. 6

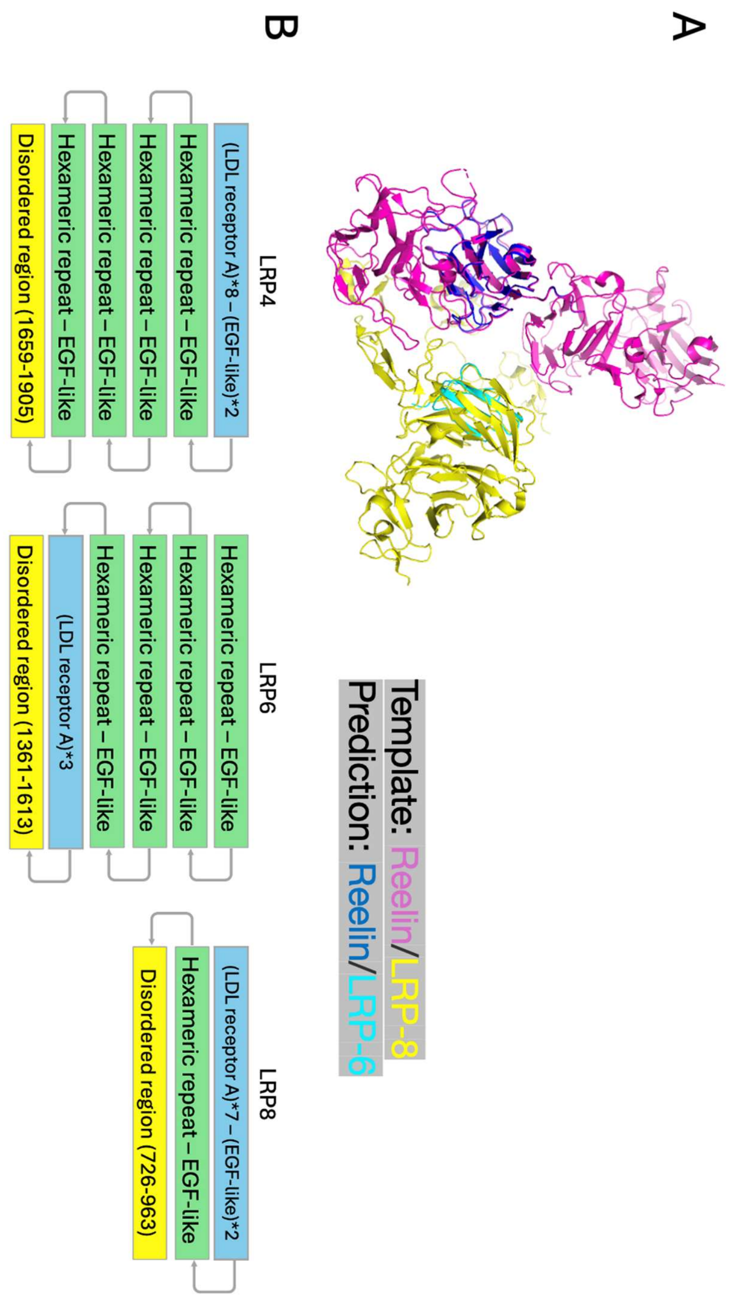


A. Predicted protein complex alignment of Reelin/Lrp-8 (canonical interaction used as template) and Reelin/Lrp-6 (predicted). B. Sequence- and structure-based comparison of LRP4, LRP6, and LRP8 proteins. All three share common structural scaffolds, but they differ in compositions.

Suppl. Fig. 7

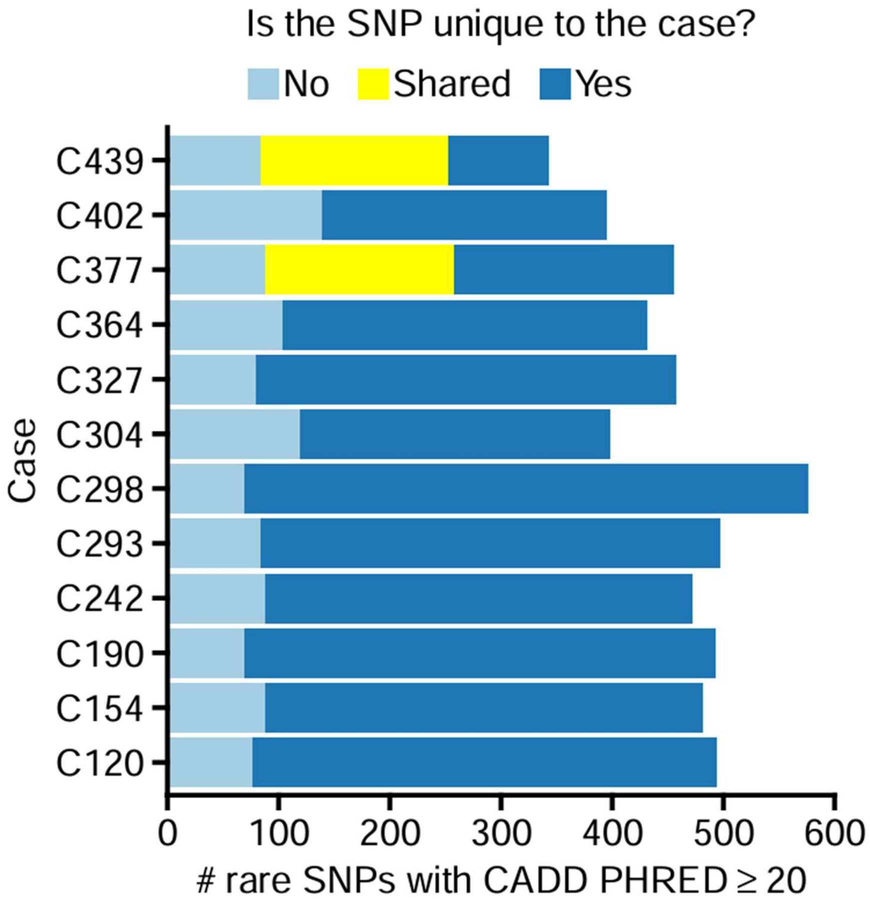


Stacked bar graph for rare gene variants number identified in ADAD and protected cases with CADD PHRED ≥ 20.

Suppl. Fig. 8

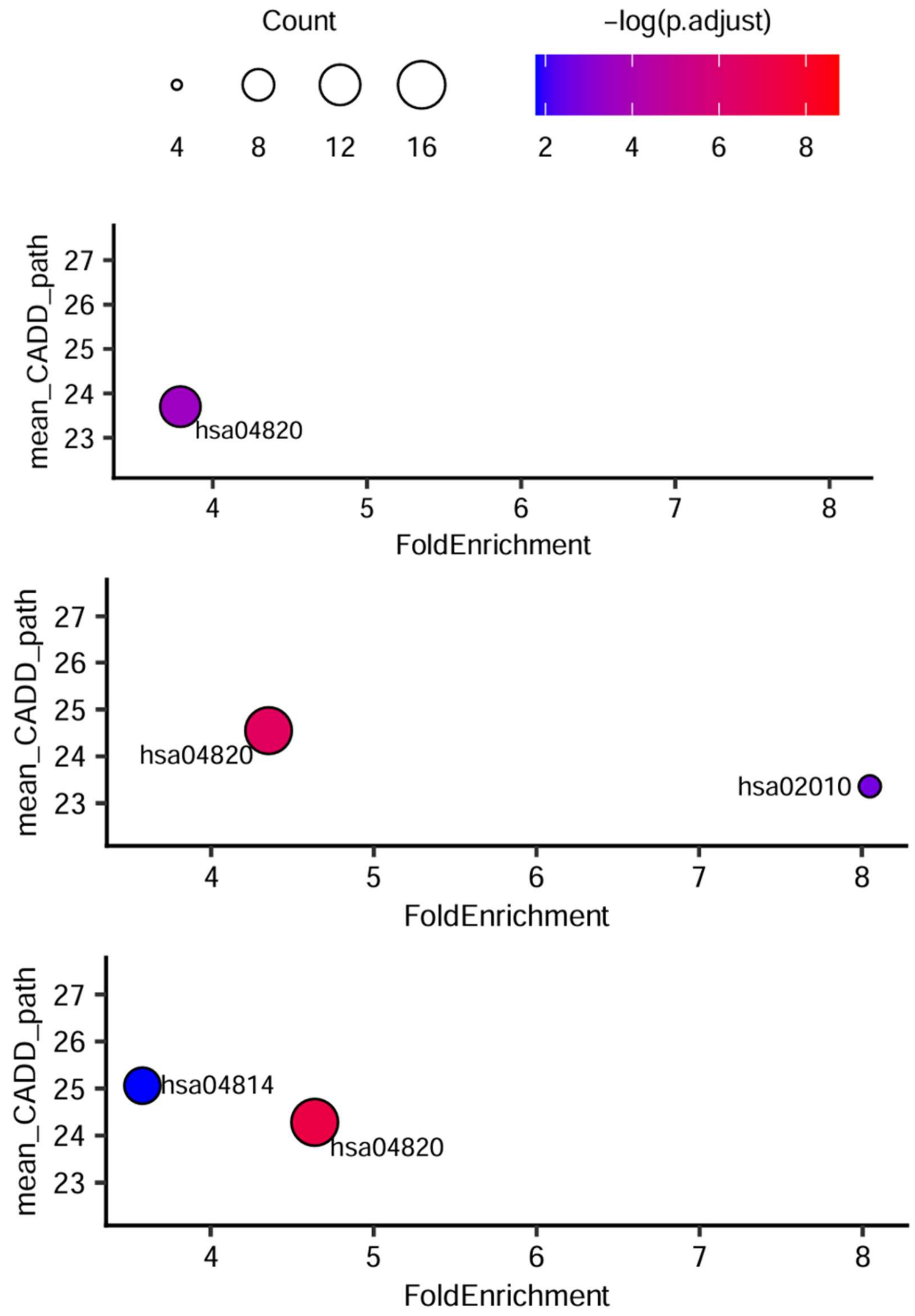


KEGG pathway enrichment for rare exonic variants found in the three ADAD cases that showed significant enrichment, plotted by enrichment ratio and average CADD (Combined Annotation Dependent Depletion) PHRED score.

Suppl. Fig. 9

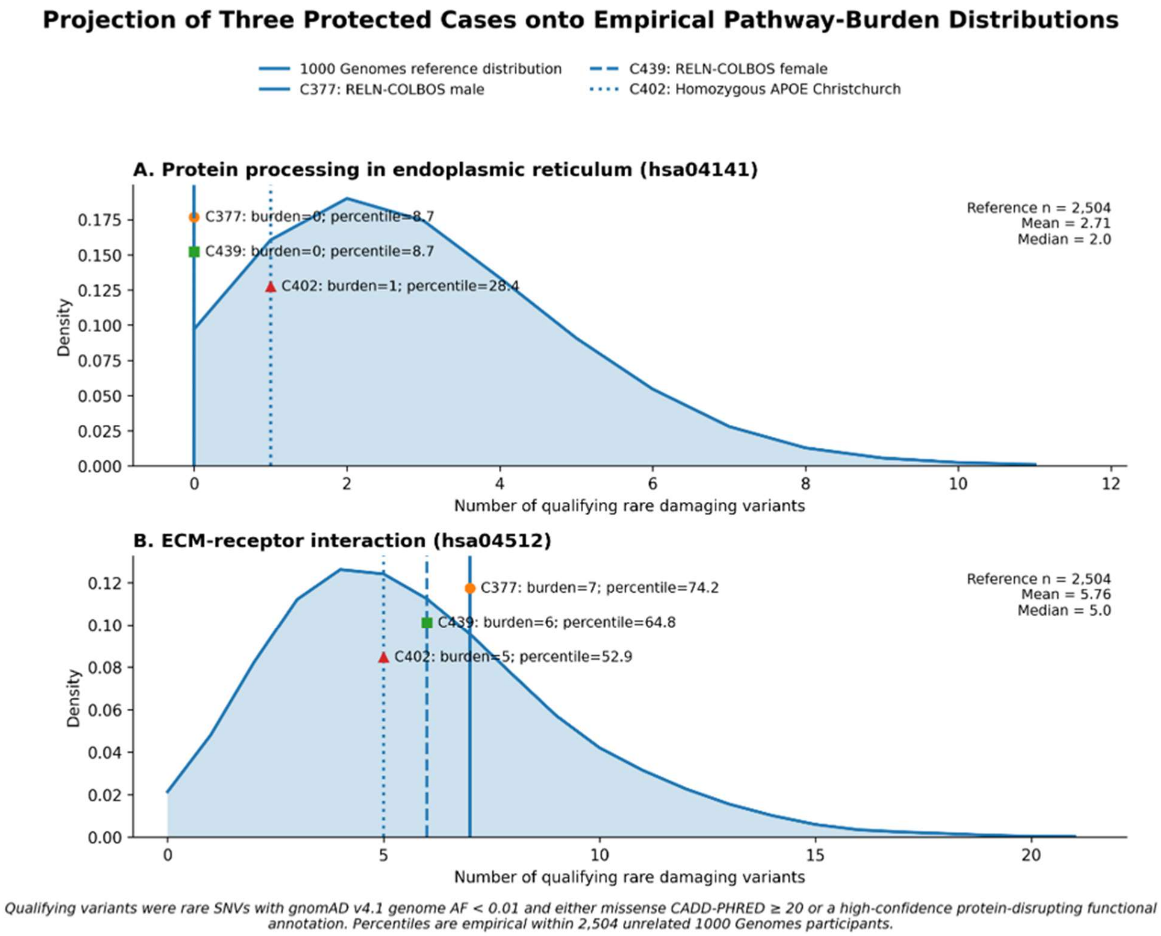


Rare exonic deleterious variants burden density plot for 1000g whole genome sequences obtained from 2504 subjects for the hsa04141 and hsa04512 pathways. The RELN-COLBOS carriers (male = orage symbol, female = green symbol) and the hoAPOECh (red symbol) carrier burden are depicted relative to general population density.

Suppl. Fig. 10

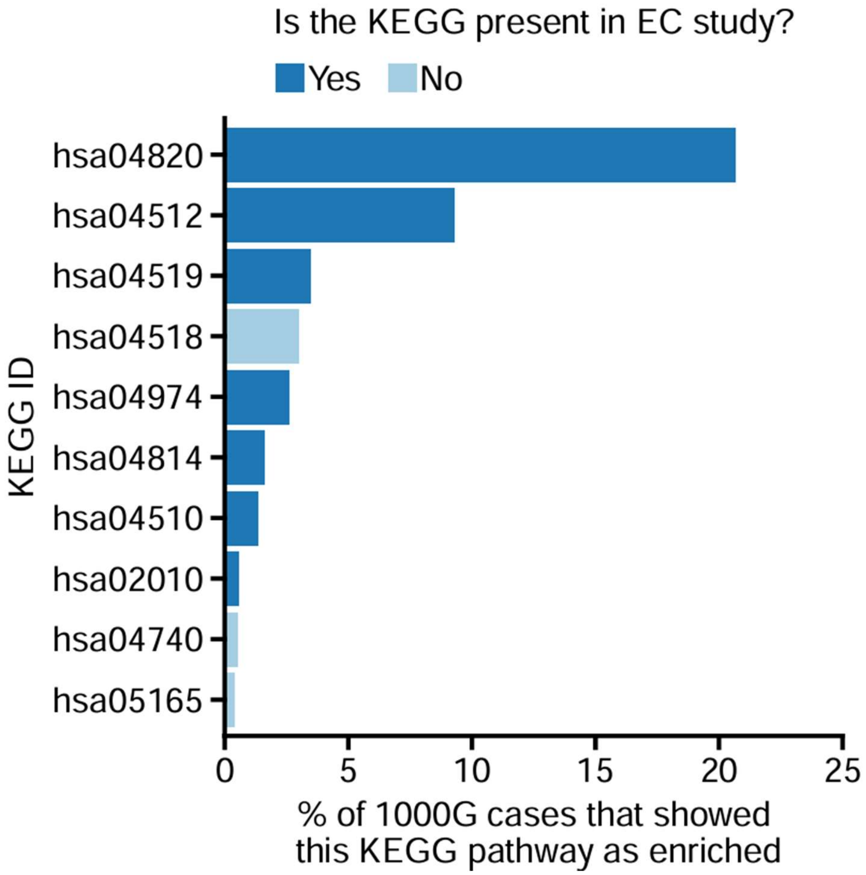


Stacked bar graph for percentage of 1000g individuals showing significant enrichment for rare exonic genes in KEGG pathways.
